# Landscape and climatic variations of the Quaternary shaped multiple secondary contacts among barn owls (*Tyto alba*) of the Western Palearctic

**DOI:** 10.1101/2021.06.09.447652

**Authors:** Tristan Cumer, Ana Paula Machado, Guillaume Dumont, Vasileios Bontzorlos, Renato Ceccherelli, Motti Charter, Klaus Dichmann, Hans-Dieter Martens, Nicolaos Kassinis, Rui Lourenço, Francesca Manzia, Laure Prévost, Marko Rakovic, Felipe Siverio, Alexandre Roulin, Jérôme Goudet

**Affiliations:** Department of Ecology and Evolution, University of Lausanne, Lausanne, Switzerland; Green Fund, Kifisia, Athens, Greece; “TYTO” - Organization for the Management and Conservation of Biodiversity in Agricultural Ecosystems, Larisa, Greece; Centro Recupero Rapaci del Mugello, Firenze, Italy; Shamir Research Institute, University of Haifa, Katzrin, Israel; Department of Geography and Environmental Sciences, University of Haifa, Haifa, Israel; Hyldehegnet 27, 6400 Sønderborg, Denmark; Gettorfer Weg 13, 24214 Neuwittenbek, Germany; Game and Fauna Service, Ministry of the Interior, Nicosia, Cyprus; MED Mediterranean Institute for Agriculture, Environment and Development, Laboratory of Ornithology, IIFA, University of Évora, Évora, Portugal; Centro di Recupero per la Fauna Selvatica–LIPU, Rome, Italy; Association C.H.E.N.E., Centre d’Hébergement et d’Etude sur la Nature et l’Environnement, 76190 Allouville-Bellefosse, France; Natural History Museum of Belgrade, Belgrade, Serbia; Canary Islands’ Ornithology and Natural History Group (GOHNIC) 38480 Buenavista del Norte, Tenerife, Canary Islands, Spain; Swiss Institute of Bioinformatics, Lausanne, Switzerland

**Keywords:** Demographic modeling, glacial refugium, Haplotypes, Population genomics, postglacial recolonization, Whole-genome resequencing

## Abstract

The combined actions of climatic variations and landscape barriers shape the history of natural populations. When organisms follow their shifting niches, obstacles in the landscape can lead to the splitting of populations, on which evolution will then act independently. When two such populations are reunited, secondary contact occurs in a broad range of admixture patterns, from narrow hybrid zones to the complete dissolution of lineages. A previous study suggested that barn owls colonized the Western Palearctic after the last glaciation in a ring-like fashion around the Mediterranean Sea, and conjectured an admixture zone in the Balkans. Here, we take advantage of whole-genome sequences of 94 individuals across the Western Palearctic to reveal the complex history of the species in the region using observational and modeling approaches. Even though our results confirm that two distinct lineages colonized the region, one in Europe and one in the Levant, they suggest that it predates the last glaciation and identify a narrow secondary contact zone between the two in Anatolia. Nonetheless, we also show that barn owls re-colonized Europe after the glaciation from two distinct glacial refugia: a western one in Iberia and an eastern one in Italy. Both glacial lineages now communicate via eastern Europe, in a wide and permeable contact zone. This complex history of populations enlightens the taxonomy of *Tyto alba* in the region, highlights the key role played by mountain ranges and large water bodies as barriers and illustrates the power of population genomics in uncovering intricate demographic patterns.

## Introduction

Species distribution patterns fluctuate in response to climatic variations, as populations relocate to follow their shifting niches^1^. When organisms colonize new areas, obstacles in the landscape may lead populations to split with varying degrees of geographic isolation. Evolution, via mutation, drift, local adaptation and gene flow, will then act independently on each of the isolated populations. If two allopatric populations are later geographically reconnected, after a certain amount of time and divergence, it can create a secondary contact zone. For example, when populations on both sides of an obstacle meet at the end of it in a ring-like fashion, as described in birds ^2^, amphibiens ^3, 4^, or plants ^5^. Likewise, climate oscillations can lead to cyclical isolation and secondary contacts between populations as regional suitability varies ^6^.

This complex interplay between climatic variations and landscape has been extensively studied in the Western Palearctic ^7, 8^, specifically in light of the cycles of glacial and interglacial periods that characterized the Quaternary ^9^. During the last glaciation, colder temperatures and the expansion of the ice sheets in the north rendered large areas unsuitable for many species, which led them to follow their niches southward. Species found refuge in the Mediterranean peninsulas and in northern Africa where climatic conditions were more amenable, forming isolated populations. At the end of the last glaciation maximum (i.e. approximatively 20k years ago ^10^), this process was reversed as the climate warmed and the melting of the continental ice caps exposed free land that could be recolonized. An extensive literature addressing the post-glacial history of European organisms describes how the complex landscape of the continent, combined with the distribution of species during the glaciation, conditioned their recolonization processes ^8,11^. However, the low-resolution of genetic data used before the genomic era was often insufficient to resolve the intricate and often fine-scale evolutionary processes that occurred during recolonization.

Rapid development of high-throughput sequencing technologies and corresponding methodological tools during the last decades has opened new avenues to study natural populations with high precision. In particular, it has allowed biologists to reconstruct the evolutionary history of species and highlight the diversity of processes acting when populations or subspecies interact in secondary contact. These processes have been found to result in a variety of situations. While the prolonged isolation of populations may lead to allopatric speciation (many examples in plants ^12,13^, amphibian ^3^, insects ^14^, mammals ^15^ and birds ^16^), secondary contact tends to show a broad range of admixture patterns. When admixture occurs, it may vary from narrow hybrid zones between lineages ^17^, to the complete dissolution of a lineage ^18^, through a gradual level of admixture along a gradient of mixing populations ^19^.

Microsatellite and mitochondrial data suggested that the barn owls (*Tyto alba*), a non-migratory raptor, colonized the Western Palearctic in a ring-like fashion around the Mediterranean Sea after the last glaciation ^20^. Under this scenario, a postglacial expansion from the glacial refugium in the Iberian Peninsula to northern Europe formed the western branch of the ring, with the eastern branch present across the Levant and Anatolia. While these observations led to conjecture of a potential admixture zone in the Balkans, the available data at the time combined with the overall low genetic differentiation in this species did not allow to fully resolve this question. Moreover, the peculiar genetic makeup of populations in the presumed contact zone brought into question the possibility of a cryptic glacial refugium in the eastern Mediterranean peninsulas.

The difficulty in resolving the post glacial expansion of barn owls is mirrored by their convoluted taxonomy in the Western Palearctic. In this region, *Tyto alba* is classified into different subspecies based on geography and plumage coloration. First, *T. a. erlangeri* (Sclater, WL, 1921) reported in Crete, Cyprus and Middle East, may match the Levant lineage. Second, *T. a. alba* (Scopoli, 1769) is white-colored and supposedly present in western Europe and western Canary Islands, and could represent the western arm of the ring colonization. *T. a. guttata* (Brehm, CL, 1831), the third subspecies is a dark rufous morph allegedly found in Northern and Eastern Europe, in the Balkans and around the Aegean Sea. This taxonomy does not match any known genetic lineage identified so far and overlaps with the area where admixture between the two lineages that colonized Europe supposedly happens, making the presence of a subspecies in this area puzzling in light of the history known so far.

Here, taking advantage of whole genome sequences of 94 individuals from all around continental Europe and the Mediterranean Sea, we elucidate the demographic history of barn owls in the Western Palearctic. Combining descriptive and modelling approaches based on genomic and ecological data, we identify how the climatic variations and landscape of the region shaped the history of this species. We also investigate how previously isolated populations of barn owls interact at secondary contacts between different lineages, and discuss the convoluted taxonomy with regards to their history.

## Material and methods

### Samples and data preparation

#### Sampling, molecular and sequencing methods

The whole genomes of 96 individual barn owls (*Tyto alba*) were used in this study (table S1): 94 individuals were sampled in 11 Western Palearctic localities: Canary Islands (Tenerife island - WC), Portugal (PT), France (FR), Switzerland (CH), Denmark (DK), Serbia (SB), Greece (GR), Italy (IT), Aegean islands (AE), Cyprus (CY) and Israel (IS). In addition, one Eastern (*Tyto javanica* from Singapore) and one American barn owl (*Tyto furcata* from California, USA) were used as outgroups. Illumina whole-genome sequences of individuals from PT, FR, CH, DK and the outgroups were obtained from the GenBank repository (BioProject PRJNA700797). For the remaining 61 individuals, we followed a similar library preparation and sequencing protocol as outlined in Machado et al. ^21^. Briefly, genomic DNA was extracted using the DNeasy Blood & Tissue kit (Qiagen, Hilden, Germany), and individually tagged. 100bp TruSeq DNA PCR-free libraries (Illumina) were prepared according to manufacturer’s instructions. Whole-genome resequencing was performed on multiplexed libraries with Illumina HiSeq 2500 PE high-throughput sequencing at the Lausanne Genomic Technologies Facility (GTF, University of Lausanne, Switzerland).

#### Data processing, SNP calling and technical filtering

The bioinformatics pipeline used to obtain analysis-ready SNPs was adapted from the Genome Analysis Toolkit (GATK) Best Practices ^22^ to a non-model organism following the developers’ instructions, as in Machado et al. ^21^. Raw reads were trimmed with Trimommatic v.0.36 ^23^ and aligned to the reference barn owl genome ^21^ with BWA-MEM v.0.7.15 ^24^. Base quality score recalibration (BQSR) was performed using high-confidence calls obtained from two independent callers – GATK’s HaplotypeCaller and GenotypeGVCF v.4.1.3 and ANGSD v.0.921 ^25^ – as a set of “true variants” in GATK v.4.1.3.

Genotype calls were filtered for analyses using a hard-filtering approach as proposed for non-model organisms, using GATK and VCFtools ^26^. Calls were removed if they presented: low individual quality per depth (QD < 5), extreme coverage (1100 > DP > 2500), mapping quality (MQ < 40 and MQ >70), extreme hetero or homozygosity (ExcessHet > 20 and InbreedingCoeff > 0.9) and high read strand bias (FS > 60 and SOR > 3). Then, we removed calls for which up to 5% of genotypes had low quality (GQ < 20) and extreme coverage (GenDP < 10 and GenDP > 40). We kept only bi-allelic sites, excluded SNPs on the heterozome (Super scaffolds 13 and 42 ^21^) and an exact Hardy-Weinberg test was used to remove sites that significantly departed (p=0.05) from the expected equilibrium using the package HardyWeinberg ^27,28^ in R ^29^, yielding a dataset of 6,448,521 SNP (mean individual coverage: 19.99X (sd: 4.38)). Lastly, we discarded singletons (minimum allelic count (mac) <2), yielding to a total of 5,151,169 SNP for the population genomic analyses.

#### SNP phasing and quality control

The set of 6,448,521 variants was phased in two steps. First, individual variants were phased using a read-based approach in which reads covering multiple heterozygous sites were used to resolve local haplotypes. To do so, WhatsHap v1.0^30^ was run independently for each individual with default parameters. Secondly, variants were statistically phased with Shape-It v4.1.2^31^. This algorithm integrates local individual phase and applies an approach based on coalescence and recombination to statistically phase haplotypes and impute missing data. Shape-It was run following the manual instructions for a better accuracy: the number of conditioning neighbors in the PBWT was set to 8, and the MCMC chain was run with 10 burn-in generations, 5 pruning iterations, each separated by 1 burn-in iteration, and 10 main iterations.

To assess the quality of the phasing, we examined phase accuracy by using the switch-error-rate metric^32^. When comparing two phasing for an individual’s variants, a switch error occurs when a heterozygous site has its phase switched relative to that of the previous heterozygous site. Thus, for each individual, we compared the *true* local phasing inferred form the read-based approach (WhatsHap) and the *statistical* phasing of this individual’s variants statistically phased by Shape-It, with read-based phase information ignored only for the individual considered (same version and parameters than in the paragraph above). The final estimation of the switch error rate was done using the switchError code to compare both phasing sets (available at https://github.com/SPG-group/switchError) (fig. S1).

### History of barn owls around the Mediterranean Sea

#### Population structure and genetic diversity

In order to investigate population structure among our samples, sNMF ^33^ was run for a number of clusters K ranging from 1 to 10 with 25 replicates for each K to infer individual clustering and admixture proportions. For this analysis, SNPs were pruned for linkage disequilibrium in PLINK v1.946 ^34^ (parameters --indep-pairwise 50 10 0.1) as recommended by the authors, yielding 594,355 SNP. Treemix ^35^ was used to calculate a drift-based tree of our populations, using this LD-pruned dataset. To detect admixture events between populations, 10 Treemix replicates were run for 0 to 10 migration events, with the tree rooted on the WC population, representative of a non-admixing population (see results, fig. S6).

Population expected and observed heterozygosity, population-specific private alleles, population-specific rare alleles (mac≤5) and population-specific total number of polymorphic sites were estimated using custom R scripts on the 5,151,169 variants dataset. To account for differences in sample sizes, which ranges from 4 to 10, population-specific statistics were calculated by randomly sampling 5 individuals from the larger populations (all except FR and SB) 10 times in a bootstrap-fashion and estimating the mean and standard deviation (SD). Individual-based relatedness (β) ^36^ and inbreeding coefficient for SNP data were calculated with the R package SNPRelate^37^. Overall F_ST_, population pairwise *F*_ST_ and population specific F_ST_^36^ were computed with the hierfstat package v.0.5-9 ^38^. Confidence intervals for population specific FST were computed by dividing the SNPs into 100 blocs, and bootstrapping 100 times 100 blocks with replacement. Finally, Principal Component Analyses (PCA) were also performed with the R package SNPRelate, first with all individuals and second only with the 66 European ones (excluding WC, CY and IS).

#### Haplotype sharing

To measure shared ancestry in the recent past between individuals, we ran fineSTRUCTURE ^39^ on the phased dataset including all individuals. For this analysis we initially modelled haplotype sharing between individuals using ChromoPainter to generate a co-ancestry matrix, which records the expected number of haplotypes chunks each individual donates to another. For this ChromoPainter step, we converted phased haps files to chromopainter phase files using the impute2chromopainter.pl script provided at http://www.paintmychromosomes.com and generated a uniform recombination map with the makeuniformrecfile.pl script. Using the version of ChromoPainter built into fineSTRUCTURE v.2.0.8, we performed 10 EM iterations to estimate the Ne and Mu parameters (switch rate and mutation rate). The model was then run using the estimated values for these parameters (respectively 570.761 for Ne and 0.0074240 for Mu), and we used default settings to paint all individuals by all others (-a 0 0). We ran fineSTRUCTURE’s MCMC model on the co-ancestry matrix for 500,000 burn-in and 500,000 sampling iterations, sampling every 10,000 iterations to determine the grouping of samples with the best posterior probability.

### Modeling of history of European barn owl

#### Maximum-likelihood demographic inference

##### Data preparation

To describe the history of barn owls in Europe, we modeled five different demographic scenarios using fastsimcoal2 ^40,41^. Given the position of Italy on the PCA (fig. 1e), its high *F*_ST_ and lower haplotype sharing with the rest of European populations (fig. 2), we tested in particular whether it could have been a cryptic glacial refugium during the last glaciation. To focus on European populations and due to computational constraints, we simplified the dataset to model the history of four populations of eight individuals (table S1): PT as representatives of the known refugium in the Iberian Peninsula ^42^; IT and GR representing the peninsulas of Italy and Balkans, respectively; and CH, a product of the recolonization of northern Europe from the Iberian refugium^42^.

**Figure 1.**
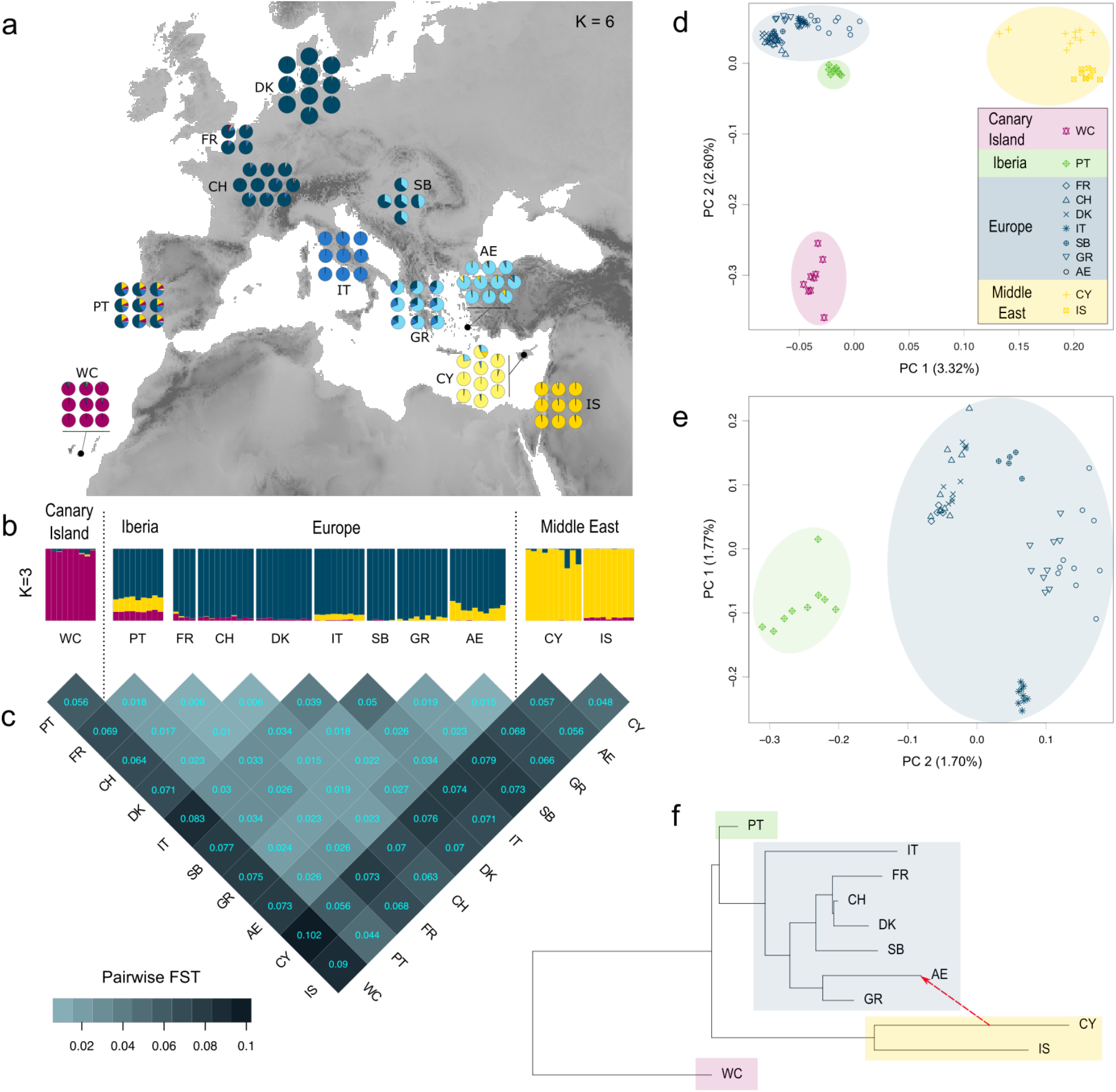
Genetic structure of barn owl populations in Western Palearctic. (**a**) Population structure for K=6. Pie charts denote the individual proportion of each of lineages as determined by sNMF and are located at the approximate centroid of the sampled population. (**b**) Population structure for K=3. Each bar denotes the individual proportion of each of the 3 lineages as determined by sNMF. (**c**) Matrix of pairwise *F*_ST_ between barn owl populations in Western Palearctic. The heatmap provides a visual representation of the *F*_ST_ values given in each cell. (**d**) PCA based on full set of 94 individuals. Point shape denotes populations and colored circles enclose sample clusters observed in sNMF (K=3). Values in parentheses indicate the percentage of variance explained by each axis. (**e**) PCA based on of the 66 European individuals. (**f**) Population tree and the first migration event in Western Palearctic populations inferred by Treemix.

**Figure 2.**
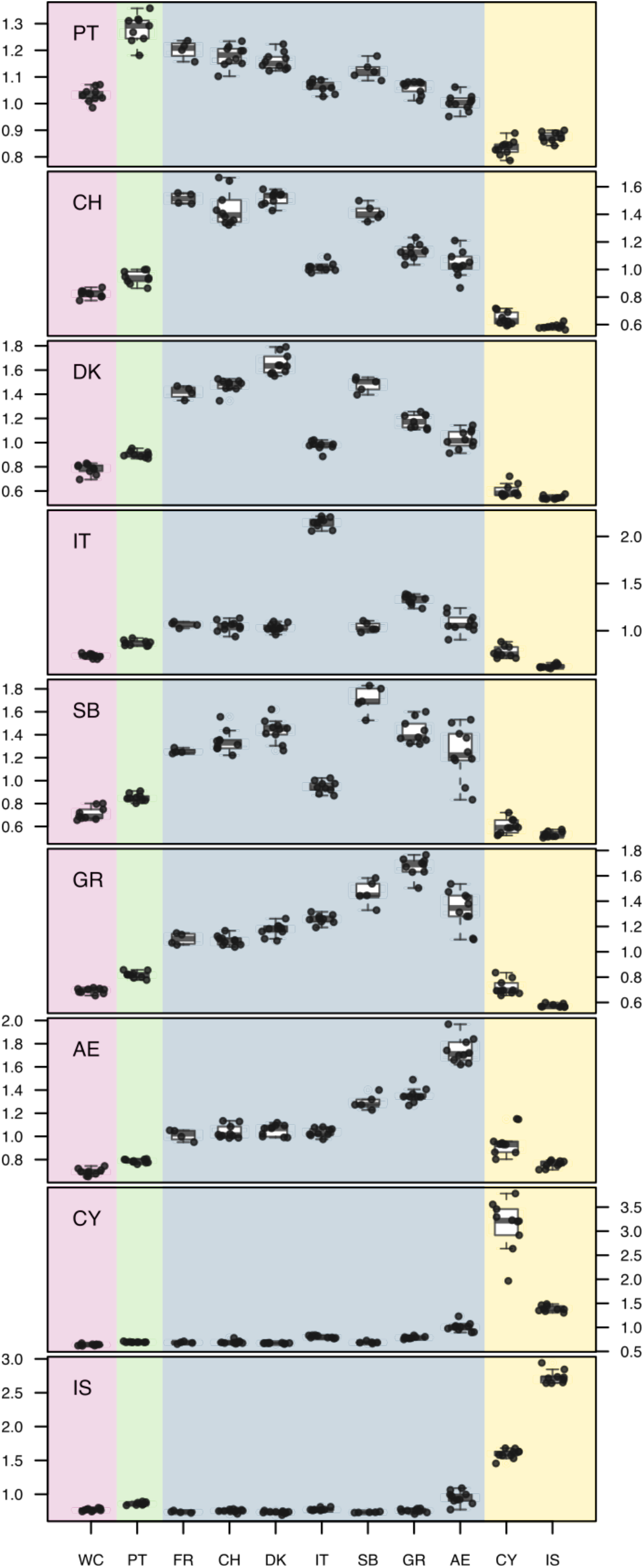
Individual haplotype sharing between barn owl populations. Part of the total length of ChromoPainter chunks inherited from other genomes. Each graph summarizes the information of all the genomes from a given population, indicated on the top-left corner. Background colors match the lineages identified in fig. 1.

Autosomal SNPs were filtered to retain only neutrally evolving regions by excluding SNPs found in genic regions and CpG mutations ^43^. To achieve homogeneity among SNPs, we removed all sites with missing data and excluded positions with a coverage outside two thirds of the standard deviation of the mean. We employed a parsimony approach based on the Tytonidae phylogenetic tree ^44^ to determine the ancestral state of the SNPs using the genomes of the two outgroups. Sites for which it was impossible to attribute a state based on the available outgroups were discarded. The remaining 770,718 SNPs were used to produce population pairwise site frequency spectra (SFS).

##### Demographic scenarios and parameters

Five different scenarios were tested to model the history of barn owls in continental Europe, with a special focus on the period since the last glaciation (fig. 2, fig. S8). Three models included only one refugium in the Iberian Peninsula and various possibilities of colonization scenarios, thus excluding the persistence of barn owls in a second refugium during the glaciation. The two last models included two refugia during the last glacial maximum (LGM), a western refugium in the Iberian Peninsula and an eastern refugium, in the Italian Peninsula. The models *1R-1, 1R-2* and *1R-3* included only one refugium in the Iberian Peninsula. *1R-1* model assumed only one colonization route around the north side of the Alps (forming the CH population), and from there move southeast to reach first the Balkans (GR) and then Italy (IT). The two other single-refugium models (*1R-2* and *1R-3*) assumed two distinct colonization routes, one north of the Alps to CH and the other south of the Alps along the Mediterranean coast to IT. *1R-2* assumes current Greece would have been colonized by owls from the Italian Peninsula, following the route along the Mediterranean coast. In *1R-3*, Greece would have been colonized via northern Europe, while the Mediterranean expansion would have stopped in Italy.

The last two models included a second, eastern, refugium (*2R-1* and *2R-2*). In these models, the western lineage expansion from the Iberian population would have colonized Europe before the last glaciation, thus occupying all the Mediterranean peninsulas. During the last glaciation, two distinct populations would have survived, respectively in the Iberian (western refugium) and Italian (eastern refugium) Peninsulas. Both models assume that northern Europe was recolonized from the Iberian lineage after the glaciation, but they differ in the scenario of recolonization of southeastern Europe. In *2R-1*, Greece was recolonized from the Italian refugium while in *2R-2* the expansion from the Iberian Peninsula would have recolonized all eastern Europe, including current Greece. In this last scenario, the Italian population would be the only relic from the eastern refugium. For all scenarios, migrations between populations was allowed (see fig. 2 and table S2).

Wide search ranges for initial simulation parameters were allowed for population sizes, divergence times and migration rates (table S2). Each population split was preceded by an instantaneous bottleneck, in which the founding population size was drawn from a log-uniform distribution between 0.01 and 0.5 proportion of current population sizes.

##### Demographic inference

Demographic simulations and parameter inference were performed under a composite-likelihood approach based on the joint SFS as implemented in fastsimcoal2 ^40,41^. For each of the five scenarios, 100 independent estimations with different initial values were run. For each run, there were 500000 coalescent simulations (option -n), with 50 expectation-maximization (EM) cycles (-M and -L). As we do not have an accurate mutation rate for barn owls, we fixed the end of the glaciation to 6000 generations BP (approximately 18’000 years BP with a 3-year generation time) and scaled all other parameters relative to it using the -0 command option (using only polymorphic sites). The best-fitting scenario out of the five tested was determined based on Akaike’s information criterion ^45^ (AIC) and confirmed through the examination of the likelihood ranges of each scenario as suggested in Kocher et al. (1989)^46^. Non-parametric bootstrapping was performed to estimate 95% confidence intervals (CI) of the inferred parameters under the best-fitting scenario. To account for LD, a block-bootstrap approach was employed as suggested by the authors ^40,41^: the SNPs were divided into 100 same-size blocks, and then 100 bootstrap SFS were generated by sampling these blocks with replacement. Due to computational constraints, for bootstrapping we ran 50 independent parameter inferences per bootstrapped SFS with only 10 EM cycles each, instead of 50 cycles used for comparing scenarios above. This procedure has been defined as conservative ^47^, and is expected to produce quite large confidence intervals. We accepted this trade-off as our main goal was to determine the best demographic topology, accepting uncertainty on specific parameter values. The highest maximum-likelihood run for each bootstrapped SFS was used to estimate 95% CI of all parameters.

#### Niche modeling

In order to identify the regions of high habitat suitability for barn owls at the last glacial maximum (LGM, 20’000 years BP) and to support the demographic scenarios tested in the previous section, we modelled the past spatial distribution of the species in the Western Palearctic. We built species distribution model (SDM) using Maximum Entropy Modelling (MaxEnt), a presence-only based tool ^48^. Current climatic variables for the Western Palearctic (fig. S10) were extracted from the WorldClim database at 5 arc min resolution using the R package rbioclim ^49^, and filtered to remove variables with a correlation of 0.8 or higher (fig. S9). The variables retained were: Mean Diurnal Range (Bio2), Min Temperature of Coldest Month (Bio6), Temperature Annual Range (Bio7), Mean Temperature of Wettest Quarter (Bio8), Precipitation Seasonality (Bio15), Precipitation of Driest Quarter (Bio17) and Precipitation of Coldest Quarter (Bio19). We built models with linear, quadratic and hinge features, and with a range (1 to 5) of regularization multipliers to determine which combination optimized the model without over complexifying it. The best combination based on the corrected AIC (as recommended by Warren & Seifert ^50^) was achieved with a quadratic model with 1 as regularization multiplier (table S5). We ran 100 independent maxent models, omitting 25% of the data during training to test the model. To avoid geographic bias due to different sampling effort in the distribution area of the species, we randomly extracted 1000 presence points within the IUCN distribution map ^51^ for each model run ^52^.

Predictive performances of the models were evaluated on the basis of the area under the curve (AUC) of the receiver operator plot of the test data. For all models with an AUC higher than 0.8 (considered a good model ^53,54^), we transformed the output of Maxent into binary maps of suitability. We assigned a cell as suitable when its mean suitability value was higher than the mean value of the 10% test presence threshold. This conservative threshold allows us to omit all regions with habitat suitability lower than the suitability values of the lowest 10% of occurrence records. Finally, we averaged the values of the models for each cell, and only cells suitable in 90% of the models were represented as such in the map.

We projected the models to the climatic conditions of the mid-Holocene (6’000 years BP) and the LGM (20’000 years BP), which we extracted from WorldClim at the same resolution as current data. When projecting to past climates, the Multivariate Environmental Similarity Surface (MESS) approach ^48^ was used to assess whether models were projected into climatic conditions different from those found in the calibration data. As our goal was to highlight only areas of high suitability for barn owls, cells with climatic conditions outside the distribution used to build the model were assigned as unsuitable (0 attributed to cell with negative MESS). For each timepoint, the results of the models were merged and transformed into a binary map as described for current data (fig. 2c).

### Barriers and corridors

#### Migration surface estimation in the western Palearctic

The Estimated Effective Migration Surface (EEMS) v.0.0.9 software ^55^ was used to visualize geographic regions with higher or lower than average levels of gene flow between barn owl populations of the Western Palearctic. Using the SNP dataset pruned for LD produced above, we calculated the matrix of genetic dissimilarities with the tool bed2diff. The free Google Maps api v.3 application available at http://www.birdtheme.org/useful/v3tool.html was used to draw the polygon outlining the study area in the Western Palearctic. EEMS was run with 1000 demes in five independent chains of 5 million MCMC iterations with a burn-in of 1 million iterations. Results were visually checked for MCMC chain convergence (fig. S11) and through the linear relation between the observed and fitted values for within- and between-demes estimates using the associated R package rEEMSplots v.0.0.1 ^55^. With the same package, we produced a map of effective migration surface by merging the five MCMC chains.

#### Isolation by distance in continental Europe

To investigate how population structure correlated with spatial distances between European populations and to detail the role of the Alps as a barrier to gene flow, we performed Mantel tests as implemented in the ade4 package v.1.7-15 ^56^ for R. We compared the genetic distances (pairwise F_ST_ between populations, see section *Population Structure and Genetic Diversity* for details) with different measures of geographical distance between populations: the shortest distance over land via direct flight and the distance constrained by the presence of the Alps, forcing the connection of the Italian population to the other populations via the Greek Peninsula (fig. S13). We also tested the linear regression between both variables in R.

## Results

### History of barn owls around the Mediterranean Sea

#### Genetic diversity and population structure in the Western Palearctic

Despite an overall low differentiation (overall *F*_ST_=0.047, comparable with the overall FST=0.045 estimated by Burri et al. ^20^), the dataset revealed a structuration of the genetic diversity among barn owls of the Western Palearctic. The first axis of the genomic PCA (explaining 3.32% of the total variance) contrasted individuals from the Levant populations (IS and CY) to all other individuals (fig. 1d), consistent with K=2 being the best estimate in sNMF (fig. S4, fig. S5). For K=3 (fig. 1b), the Canary population (WC) formed an independent genetic cluster, and this was confirmed by the second axis of the PCA (explaining 2.6% of the variance) opposing it to all other individuals (fig. 1d, fig. S2). This isolation of WC was also observable in table 1, with a lot of privates and rare alleles in Tenerife island, it’s higher F_IT_ and the highest population specific F_ST_ of all sampled populations. On the same PCA (fig. 1d), individuals from European populations (FR, CH, DK, IT, SB, GR, AE) formed a third distinct cluster, matching their grouping in a single cluster at K=3 with sNMF (fig. 1b). The Iberian individuals (PT) occupied a central position on the PCA (around 0 on both axes) and a mixed composition in sNMF (fig. 1b). This central position of the Iberian population was also visible in the pairwise *F*_ST_, where the highest value in the pairs involving PT is 0.055 (with both CY and WC) while all other pairwise comparisons involving populations from two of the distinct groups identified before (Levant, Canary Islands and Europe) have values equal or higher (fig. 1c). The TreeMix analysis (fig. 1f) was also consistent with these results, since it also identified these major lineages, first isolating the Canary population in a specific lineage, then grouping the Levant populations (IS and CY) in a second lineage and finally all European populations in a third lineage. In Europe, the Iberian population was basal to all other populations.

**Table 1.** Population genetic diversity, inbreeding and divergence estimates for 11 populations of barn owls from the Western Palearctic. Standard deviations of the mean are provided between brackets for each parameter, see Methods section for details.

| *Pop* | *N* | *#PI* | *#PA* | *#Rare* | *F<sub>IT</sub>* | *F<sub>IS</sub>* | *Pop F<sub>ST</sub>* |
| --- | --- | --- | --- | --- | --- | --- | --- |
| *WC* | 9 | 2217235<br>(37007) | 123016<br>(2465) | 407403<br>(10568) | 0.067 (0.054) | -0.022 (0.057) | 0.116 (0.006) |
| *PT* | 9 | 2639343<br>(22857) | 102252<br>(2305) | 539157<br>(10106) | -0.018 (0.042) | -0.008 (0.042) | 0.003 (0.003) |
| *FR* | 4 | 2151627<br>(0) | 30290<br>(581) | 287746<br>(1468) | 0.039 (0.124) | 0.043 (0.131) | 0.050 (0.005) |
| *CH* | 10 | 2494462<br>(10723) | 47532<br>(730) | 386654<br>(3117) | 0.025 (0.019) | -0.011 (0.019) | 0.036 (0.002) |
| *DK* | 10 | 2410615<br>(15558) | 40650<br>(1378) | 349800<br>(5434) | 0.026 (0.020) | -0.02 (0.021) | 0.049 (0.002) |
| *IT* | 9 | 2404069<br>(7267) | 66297<br>(1638) | 401842<br>(4483) | 0.035 (0.012) | -0.022 (0.012) | 0.052 (0.005) |
| *SB* | 5 | 2336060<br>(0) | 32096<br>(1515) | 326906<br>(2519) | 0.025 (0.011) | -0.038 (0.011) | 0.056 (0.004) |
| *GR* | 9 | 2454653<br>(7365) | 44996<br>(1146) | 378422<br>(4152) | 0.018 (0.028) | -0.016 (0.027) | 0.039 (0.002) |
| *AE* | 10 | 2460422<br>(20650) | 56260<br>(3439) | 403259<br>(10465) | 0.018 (0.060) | -0.001 (0.062) | 0.030 (0.002) |
| *CY* | 10 | 2338318<br>(48463) | 113377<br>(2229) | 480021<br>(13581) | 0.022 (0.047) | -0.034 (0.049) | 0.059 (0.004) |
| *IS* | 9 | 2509099<br>(9500) | 172624<br>(4018) | 608944<br>(3597) | -0.019 (0.015) | -0.036 (0.016) | 0.021 (0.003) |

N: number of individuals in the population; #PI: number of polymorphic sites per populations; #PA: number of private alleles per population; #Rare: Number of rare alleles per population; $F_{IT}$ : mean individual inbreeding coefficient relative to the meta-population; $F_{IS}$ : population level inbreeding coefficient; $F_{ST}$ : population specific $F_{ST}$ as in Weir and Goudet 2017. Populations: WC – Canary Islands, PT – Portugal, FR – France, CH – Switzerland, DK – Denmark, IT – Italy, SB – Serbia, GR – Greece, AE – Aegean Islands, CY – Cyprus, IS – Israel.

#### Population structure in continental Europe

Focusing only on European samples, southern populations (PT, IT, GR and AE) harbored a higher genetic diversity than northern populations (CH, FR, DK, SB) (table 1). The first axis of the genomic PCA based on samples from European-only populations opposed the Italian individuals to all others (fig.1e, fig. S3). This isolation of Italian samples was also apparent in the pairwise *F*_ST_ within Europe, with all the largest values involving IT. These results were consistent with sNMF on all samples for K higher than 3, where IT individuals formed an independent genetic cluster (fig. S5, illustrated in fig. 1a), as well as TreeMix, where the Italian population was the first to split and had the longest branch within European lineage (fig. 1f). Consistently with the distribution of the individuals in the European PCA (fig. 1e, fig. S3), the ancestry coefficients in the sNMF analyze of k=4 and k=5 (fig. S5) revealed the genetic differentiation of northern populations (FR, CH and DK) compared to the Italian one, also opposed in the first axis. Individuals from GR and AE individuals shared ancestry with both western and Italian population, in line with their central position along this first axis of the European PCA. For k=5, a third European component was distinguished in the Aegean individuals. This component was the majoritarian in Greek samples with contributions of both northern and Italian component; and Serbian samples appeared as a mix of the northern and the Aegean component. The eastern lineage (CY and IS), grouped at previous K, were split into two distinct ancestry pools at K=6 (fig. 1a). AE individuals harbored low amounts of CY ancestry, absent in all other European populations, and two CY individuals carried a large contribution of the Aegean component. Consistently, the first migration event detected by TreeMix was from CY, a population from the Levant lineage, to AE, a population from the European lineage (fig. 1f).

#### Fine Structure and Haplotype sharing

The clustering of individuals by FineStructure, based on shared haplotypes between individuals was consistent with previous results (fig. S7). Individuals from the different populations sampled were monophyletic, except for CH and FR individuals, mixed in the same population. Consistently with this grouping, haplotypes from any given population were more likely to be found in individuals from the same population, followed by its most related populations (fig. 2). In the Levant lineage, IS haplotypes mostly painted IS Individuals but also CY individuals and vice versa. Iberian haplotypes mostly painted PT individuals, but also contributed greatly to the painting of all European individuals, decreasing with distance. Western European haplotypes (from FR and CH) mostly painted western European individuals, then northern individuals (from DK) and finally eastern individuals (from SB, GR and AE). The reverse pattern was observed for haplotypes from eastern Europe (GR, AE), with a gradient of contribution decreasing from east to west. Haplotypes from DK and SB mostly painted individuals from their own population, but also in their respective neighbors in both eastern and western European populations. Italian haplotypes were the most distinct haplotypes among European populations, mostly painting Italian individuals, followed by Greek individuals, and being painted by other populations at a lower rate than expected given its geographic position. Finally, AE haplotypes also painted more often CY individuals than IS individuals. This painting of levant individuals by AE haplotypes was higher than the contribution from any other European individual, and both CY and IS haplotypes painted more AE individuals than any other European individual.

### Modeling of the history of European barn owls

#### Species distribution modeling

Habitat suitability projections showed that, from a climatic point of view, there were suitable regions for barn owls all around the Mediterranean Sea during the glaciation (20’000 years BP; fig. 3c). Large areas were suitable in northern Africa and the Iberian Peninsula, but also in the two eastern Mediterranean peninsulas (current Italy and Greece). At this point, the sea levels were lower than today’s and the two eastern peninsulas were more connected, allowing for a continuous region of suitable barn owl habitat. At the mid-Holocene (6’000 years BP), major changes in sea level revealed a coastline very similar to nowadays. Our projections revealed a reduction of habitat suitability in northern Africa at this time, while the suitability of western and northern Europe increased (fig. 3c). Finally, today, nearly all continental Europe is suitable for the barn owls, with the notable exception of mountain areas (fig. 3c).

**Figure 3.**
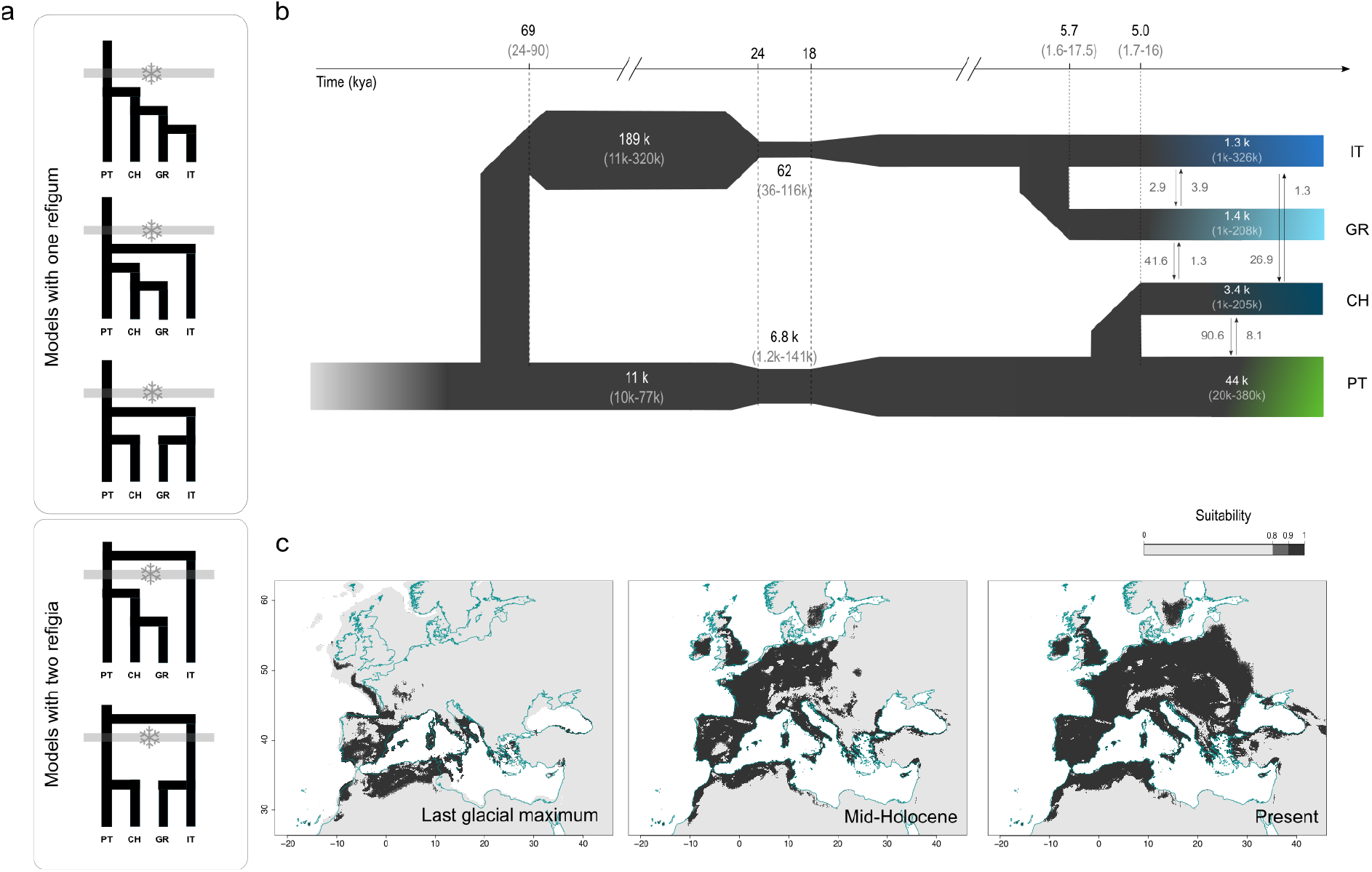
Modeling of the history of the barn owl in Europe. (**a**) Schematic representation of the five demographic scenarios tested for the colonization of the Europe by barn owls. Three models included one refugium in the Iberia during the LGM while the last two included two refugia, one in Iberia and the second in Italy. Grey bars with snowflakes represent the last glaciation. (**b**) Best supported demographic model for the history of European barn owl populations as determined by fastsimcoal2. Time is indicated in thousands of years, determined using a 3-year generation time, confidence intervals at 95% are given between brackets. Population sizes (haploid) are shown inside each population bar; arrows indicate forward-in-time migration rate and direction. (**c**) Species distribution model of barn owls based on climatic variables, projected into the past (last glacial maximum (20 kya) mid-Holocene (6 kya)) and today’s condition. Locations in dark grey were highly suitable in 90% of the models. Below that threshold cells were considered as unsuitable (lightest grey shade on the graph). The present coastline is outlined in blue in all graphs.

#### Demographic inference

AIC and raw likelihood comparisons showed that the two refugia model *2R-1* explains best the SFS of our dataset (table S3; Fig. 3b). In this model, an ancestral Italian lineage (IT) split from the Iberian lineage (PT) before the last glaciation, estimated at approximately 69’000 years BP (95% CI: 24’000-90’000 years BP; calculated with 3-year generation time). After its initial expansion, the ancestral population is estimated to have been larger in the Italian Peninsula that in Iberia (respectively 189K (11K-320k) and 11k (10K-73k) haploid individuals). During the glaciation (fixed between 24 and 18K years BP), both populations experienced a bottleneck, with a population size reduced to 6.8k (1.2k-141k) individuals in the Iberian lineage and 62 (36-116k) in the Italian. After the glaciation, the size of both populations increased to their current size, estimated at 44k (20K-380k) in the Iberian Peninsula and 1.3k (1k-326k) in the Italian Peninsula, both smaller but consistent with their estimated census size (55k-98k ^57,58^ and 6k-13k ^59^, respectively). The Greek population split from the Italian branch around 5’700 years BP while the Swiss (CH) population split slightly later from Iberia (5’000 years BP) and maintain a high level of gene flow (estimated to 90 (46-1.4k) from CH to PT and 8 (3-62) in the reverse direction). Current effective population sizes of the CH and GR populations are estimated to 3.4k (1k-205k) and 1.4k (1k-208k), respectively (fig. 2b), in line with census results (1000-2500 ^60^ and 3000-6000 ^61^, respectively). Migration between these populations is estimated to be highest from IT and GR to CH (respectively 27 (0.2-157) migrants from IT and 42 (0.1-156) from GR) and lowest in the opposite direction (respectively 1.3 (0.1-96) migrants from CH to GR and 0.02 (0.2-38) from CH to IT) (table S4). Point estimates with 95% confidence intervals for all parameters of the best model are given in (table S4), as well as single point estimates for all models (table S2).

### Barriers and corridors

#### Migration Surface Estimate in the Western Palearctic

Estimated Effective Migration Surface identified large water bodies, especially in the eastern Mediterranean and around Cyprus, as regions resisting to migration (fig. S12). On the mainland, barriers to gene flow matched the main formations of the Alpide belt in the region, an orogenic formation spanning from western Europe to eastern Asia. From west to east, a light barrier overlapped with the Pyrenees, a strong barrier spanned the Alps to the Balkans and a third obstacle matched the Taurus mountains in Anatolia. A region with high gene flow was identified in continental Europe above the Alps, spanning from western Europe to the Balkan Peninsula.

#### Isolation by distance in Europe

In continental Europe, the shortest path overland did not correlate significantly with genetic distance (fig. S13) (mantel test, p-value = 0.193, R = 0.20 / linear model, p-value = 0.26, R^2^ = 0.012). On the contrary, when the geographic distance between populations included the barrier formed by the Alps (i.e. the Italian population was connected to other populations via the Greek Peninsula, itself connected to western Europe via northern Europe), both tests were significant (mantel test, p-value = 0.002, R = 0.68 / linear model, p-value = 1.3×10^−5^, R^2^ = 0.507).

## Discussion

The history of natural populations is shaped by the combination of landscape barriers and climatic variations that isolate and mix lineages through their combined actions. Consistently with previous work ^20^, we show that barn owls colonized the Western Palearctic in a ring-like fashion around the Mediterranean Sea, with one arm around the Levant and the second throughout Europe. However, using whole genome sequences we found this colonization actually predates the last glaciation and pinpoint a narrow secondary contact zone between the two lineages in Anatolia rather than in the Balkans. In addition, we provide evidence that barn owls recolonized Europe after the LGM from two distinct glacial refugia – a western one in Iberia and an eastern in Italy – rather than a single one as it was previously thought. As temperatures started rising, western and northern Europe were colonized by owls from the Iberian Peninsula while, in the meantime, the eastern refugium population of Italy had spread to the Balkans (fig. 4). The western and eastern glacial populations finally met in eastern Europe. This complex history of populations questions the taxonomy of the multiple *Tyto alba* subspecies, highlights the key roles of mountain ranges and large water bodies as barriers to gene flow for a widespread bird and illustrates the power of population genomics in unraveling intricate patterns.

**Figure 4.**
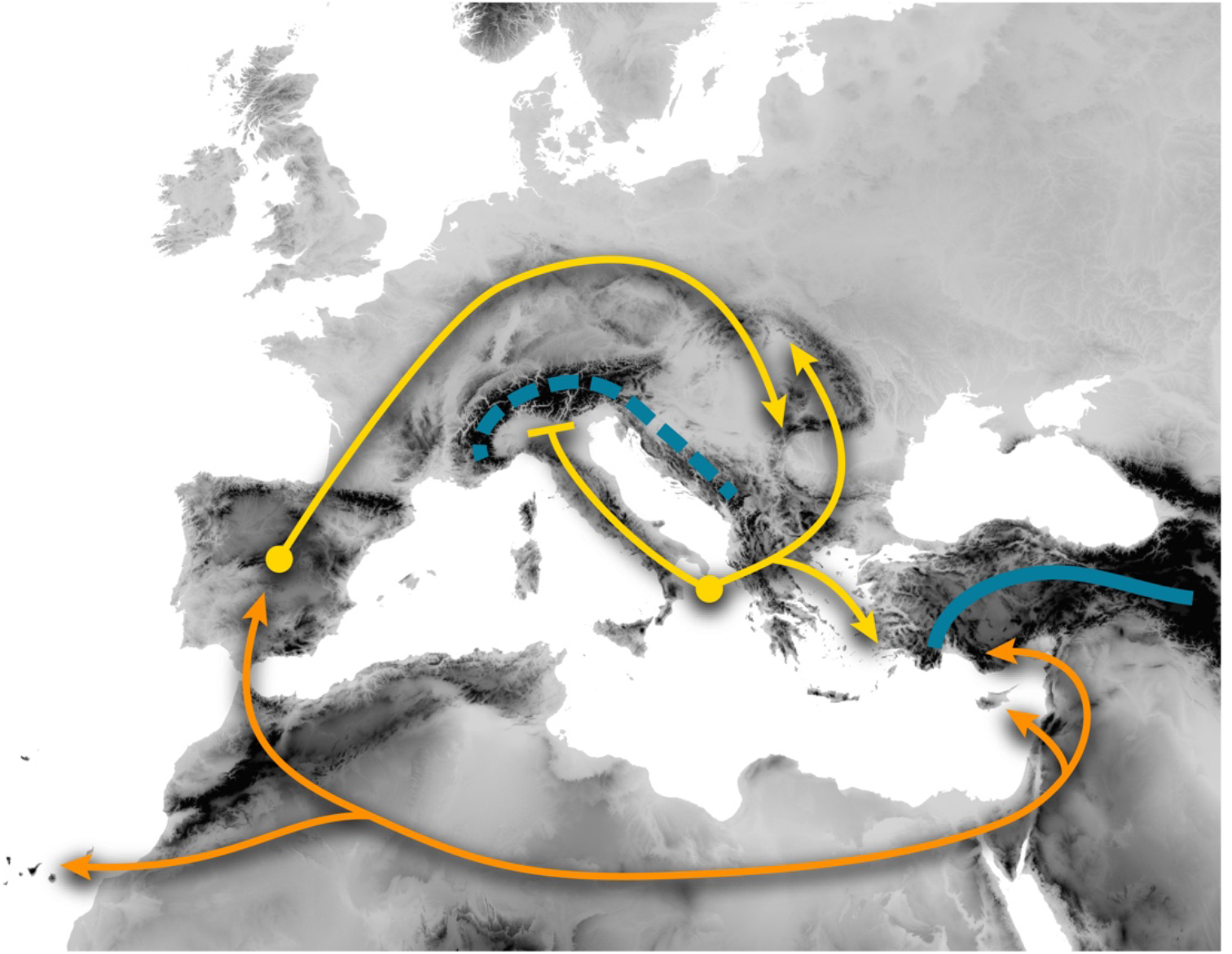
Schematic representation of the history of barn owls in the Western Palearctic and the main barriers in the region. Orange arrows depict the colonization of the region by the three main lineages (Levant, Canary Islands, European). Yellow arrows represent the modeled postglacial recolonization scheme of Europe, with two distinct refugia (yellow dots). Blue lines represent the main barriers identified in this work, namely the Alps in Europe (dashed line) and the Taurus and Zagros mountains in Anatolia (solid line).

### Colonization of the Western Palearctic and gene flow in Anatolia

Our results show that two distinct barn owl genetic lineages surround the Mediterranean Basin: one in the Levant, and a second in Europe (fig. 1b, 1c, 1d, 1f), likely connected via northern Africa. Supported by the higher and specific diversity of the basal population of each arm (namely, IS and PT; table 1), these observations are consistent with the ring colonization scenario hypothesized by Burri et al. ^20^. However, we show that a barn owl population survived the last glaciation in Italy (see next section for details) and that its genetic makeup resembles the European lineage (fig. 1b, 1d, 1f). Therefore, the ring colonization of Europe around the Mediterranean appears to have been pre-glacial, whereas the post-glacial history is more convoluted (see next section).

In previous studies, the ancestry of Greek and Aegean populations was unclear, with a hypothesized mixed origin between the European and Levant lineages ^20^. This uncertainty was mostly likely due to the low resolution of genetic markers (Mitochondrial DNA and microsatellites), as the genomic data reported here clearly show that Greek and Aegean owls are genetically much closer to European than to Levant ones (fig. 1b, 1c, 1d, 1f). This observation indicates that the European lineage reached further east than previously assumed, allowing us to pinpoint the secondary contact zone between the European and Levant lineages to Anatolia, instead of the Balkans as it had been proposed. In Anatolia, the Taurus and Zagros mountain ranges form an imposing barrier that appears to have stopped the expansion of the Levant lineage both during the ring colonization and nowadays.

Despite the barrier, and although we do not see a complete admixture of the two lineages, there is evidence of some gene flow. Indeed, the first migration in Treemix (fig. 1f) pointed to a secondary contact between CY and AE, consistent with the signals of admixture between these populations (fig. 1a, 1b, 2). The admixture pattern however is restricted geographically to this narrow region and does not permeate further into either of the lineages, as surrounding populations (IS in the Levant and GR and SB in Europe) do not show signals of admixture (fig. 1, 2). Thus, the migration between populations on both sides seems limited and possibly only occurs along a narrow corridor along the Turkish coast where only a few barn owls have been recorded ^62^. Further analyses with samples from Anatolia should allow to characterize in high resolution how and when admixture occurred in this region.

### Glacial refugia and recolonization of Europe

Previous studies showed that barn owls survived the last glaciation by taking refuge in the Iberian Peninsula and maybe even in emerged land in the Bay of Biscay ^21,42^. The observed distribution of diversity in Europe, and especially the specific makeup of the Italian populations, is best explained by a demographic model with two glacial refugia – one in Iberia and a second in Italy, derived from the Iberian population before the glaciation (fig. 3a and 3b; table S5). Environmental projections not only support this model, as Italy was highly suitable for the species at the time, but also show that, due to the low levels of the Adriatic Sea, the suitable surface extended to the west coast of the Balkans (LGM - fig. 3c). Crucially, three of the barn owl’s key prey also had glacial refugia in the Italian and Balkan peninsulas, namely the common vole (*Microtus* sp.) ^63^, the wood mouse (*Apodemus* sp.) ^64^ and shrews (*Crocidura* sp.) ^65^. The inferred size of the pre-glacial population of barn owls inhabiting current Italy was larger than any other (189k (11K-320K)), prior to a strong bottleneck during the glaciation (population reduced to 62 individuals (36-116k)). These values are likely inflated by necessary simplification of the model (e.g. instant bottlenecks) or by gene flow from unmodeled/unsampled populations, for example from the Levant lineage to eastern European populations (GR), or from northern Africa to Italy via Sicily. The latter would have been facilitated by the increased connectivity between Italy and North Africa during the glaciation (fig. 3c) ^66,67,68^.

With the warming following the LGM, Europe became gradually more suitable and, by the mid-Holocene (6000 years ago), most of western and northern Europe were appropriate for barn owls (fig. 2b) as well as the common vole (*Microtus arvalis*)^69^. The genetic similarity between Iberian (PT) and northwestern populations (CH, FR, DK; fig. 1a, 1e) indicates that barn owls colonized these newly available regions from the Iberian refugium as previously thought ^42^. The contribution from the Italian refugium to northern populations appears to have been hindered by the Alps (see next sections), as suggested by the higher genetic distance between them (fig. 1a, 1c, 1d and 2). Instead, at this time the Adriatic Sea had neared today’s levels, isolating genetically and geographically the Italian refugium from its component in the Balkan Peninsula (fig. 3b – IT-GR split ^~^6k fig. 3b and fig. 3c). Only more recently did the rise of temperatures allow for areas in the east of Europe to become suitable, finally connecting the southeastern populations near the Aegean Sea (GR and AE) with populations in the northeastern part of Europe (fig. 3c). In particular, the high heterozygosity and admixed ancestry of Serbian individuals (table 1, fig. 1a) suggest that the suture between the Iberian and the Italo-Greek glacial lineages took place in eastern Europe. This newly identified postglacial recolonization scheme of continental Europe by the barn owl matches the general pattern described for the brown bear (*Ursus arctos*) and shrew (*Sorex* sp.) ^7^.

The isolation by distance pattern observed between European populations (fig. S13) highlights a diffusion of alleles in the European populations rather than a narrow hybrid zone. Further, the inferred migration rates support high current gene flow in the region (fig. 3b, CH and GR in table S4), and we found signals of each ancestry in populations far from the suture zone (dark blue in GR and light blue in CH and DK, fig. 1a). Finally, the measure of haplotype sharing decreases consistently with distance between populations around the northern side of the Alps between populations from Iberia to Greece, excluding IT (fig. 2). Surrounded by the sea and the Alps, Italy is the exception and appears to have avoided incoming gene flow (fig. S12, isolation in fig. 2), thus being a better-preserved relic of the refugium population (own cluster in sNMF K>4, fig. S5). In contrast, the Balkan component admixes smoothly with the other European populations. Such seamless mixing of the two glacial lineages from southern refugia where barn owls are mostly white ^20,70^, brings further into question the subspecies *T. a. guttata*.

### The case of Tyto alba guttata

Traditionally, in Europe, the eastern barn owl (*T. a. guttata* (Brehm, CL, 1831)) is defined by its dark rufous ventral plumage in contrast to the white western barn owl (*T. a. alba* (Scopoli, 1769)) ^71^. With a wide distribution, it is recorded from The Netherlands to Greece, including most of northern and eastern Europe ^72^. However, this repartition does not match the history of any specific glacial lineage identified above, nor any genetically differentiated population. The dark populations of northern Europe (DK) are genetically as similar to lighter western populations (FR, CH) than to the dark ones in the east (SB). This color variance within European populations has been shown to be maintained through local adaptation ^70^, and while the genomic basis and history of this trait remain worthy of future investigations, we suggest that all European barn owls form a single subspecies (*T. a. alba*), reflecting the entire European population, regardless of their color.

### Barriers and corridors shape the connectivity of the Western Palearctic meta-population

The partition of genetic diversity among barn owls in the Western Palearctic allowed us to identify barriers and corridors to gene flow. Populations isolated by large water bodies have accumulated substantial genetic differences as, for example, the higher *F*_ST_ in the Canary and Cyprus islands (fig. 1c, table 1), and reflect the importance of water as a barrier to dispersion in this species ^21^. On the mainland, and as described for the American barn owl ^73^, major mountain ranges act as significant obstacles to migration for European barn owls and can generate genetic structure. First, the high mountain ranges of Taurus and Zagros coincide with the contact zone between the Levant and the European lineages both nowadays and potentially at the time of the pre-glacial ring colonization of Europe (see above). Second, the Alps and the Balkan Mountains slowed the northward expansion of the glacial populations of Italy and Greece after the LGM and still constrain migration between populations on both sides of their ranges. If these results remain to be confirmed with observational data (i.e. ringing data not available for all countries), they emphasize that, despite its worldwide repartition and its presence on many islands, the connectivity of barn owl populations is heavily driven by biogeographical barriers.

## Conclusion

The combination of whole genome sequencing and sophisticated modeling methods revealed the complex history of the barn owl in the Western Palearctic with a precision previously unachievable. It allowed the localization of a secondary contact zone as well as the discovery of a cryptic glacial refugium. However, several questions remain unanswered, awaiting for relevant samples to be collected and analyzed: What role did northern African populations played in connecting the Levant and European lineages? Did they contribute to the diversity observed in Italy? How narrow is the contact zone between the Levant and European lineages in Anatolia? Lastly, the origin of barn owls from the Western Palearctic as a whole also deserves further investigation, as they are believed to have colonized the Western Palearctic from the east, given the species’ supposed origin in southeastern Asia approximately 4 million years ago. But this is at odds with the higher genetic diversity of the Iberian population compared to the Levant one. Such inconsistency points to the need for samples from around the world, to understand how this charismatic group of nocturnal predators conquered the entire planet.

Research on postglacial recolonization and the subsequent phylogeographic patterns peaked at the turn of the century, with many studies providing an overview of the history of a wide variety of organisms (reviewed by Hewitt^7^). The rise in availability of genomic data for non-model species, combined with the type of approaches used here, will rewrite the history of many of them. Furthermore, it will allow to detail the genomic consequences of such history both from a neutral and selective perspective. Applied to several species, these approaches will redefine with greater clarity the broad phylogeographical patterns in the Western Palearctic and elsewhere, to re-think taxonomic classifications and to better understand how organisms might adapt to a changing environment in a complex, fragmented and rapidly changing landscape.

## Acknowledgements

We thank the following institutions and individuals for providing samples or aiding in sampling: The European Barn Owl Network, Sylvain Antoniazza and Reto Burri, the Natural History Museum of Tenerife, the Wildlife Rehabilitation Centre La Tahonilla (Cabildo Insular de Tenerife), Kristijan Ovari and the Palić Zoo (Serbia). We thank Céline Simon and Luis San-José for their valuable assistance with molecular work as well as Alexandros Topaloudis for scripts. Finally, we also thank Olivier Delaneau for his advice on the phasing of the individuals and its evaluation. This study was funded by the Swiss National Science Foundation with grants 31003A-138180 & 31003A_179358 to JG and 31003A_173178 to AR.

## Data Accessibility

The raw Illumina reads for the whole-genome sequenced individuals are available in BioProject PRJNA700797 and BioProject PRJNA727977.

## Author Contribution

TC, APM, AR, JG designed this study; GD and APM produced whole-genome resequencing libraries and called the variants; TC and APM conducted the analyses; KD, RL, JL, HDM, PB, VB, MC, KD, HDM, NK, RL, FM, KO, LP, MR and FS provided samples to the study; TC led the writing of the manuscript with input from APM and all authors.

