## Supplementary material for "Landscape and climatic variations of the Quaternary shaped multiple secondary contacts among barn owls (*Tyto alba*) of the Western Palearctic": Table and Figures Supp

*Supplementary Materials*

**Samples and data preparation**

Figure S1 ……………………………………………………………………………………………………………….. 2

**History of barn owls around the Mediterranean Sea**

*Haplotype sharing*

Figure S7 …………………………………………………………………………………………………………..…… 7

**Modeling the History of European barn owl**

*Maximum-likelihood demographic inference*

Figure S8 ……………………………………………………………………………………………………………….. 8

*Table S2* …………………………………………………………………………………………………………………. 9

*Table S3* ………………………………………………………………………………………………………………… 10

*Table S4* ………………………………………………………………………………………………………………… 11

*Niche modeling*

Figure S9 ………………………………………………………………………………………………………………. 12

Figure S10 …………………………………………………………………………………………………………….. 13

Table S5 ………………………………………………………………………………………………………………… 13

**Barriers and corridors**

*Migration Surface Estimate*

Figure S11 …………………………………………………………………………………………..………………… 14

Figure S12 ……………………………………………………………………………………………..……………… 15

*Isolation by distance*

Figure S13 ………………………………………………………………………………………………..…………… 16

**Samples and data preparation**

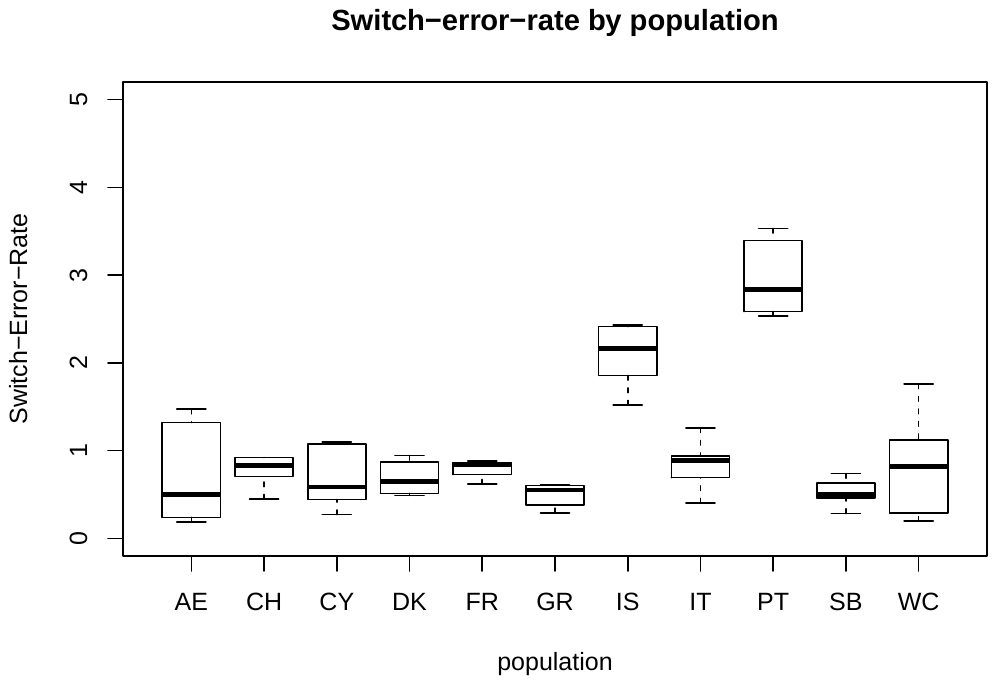

**Figure S1**– Individual phasing switch error rate (in %) grouped per population.

**History of barn owls around the Mediterranean Sea**

*Population Structure and Genetic Diversity*

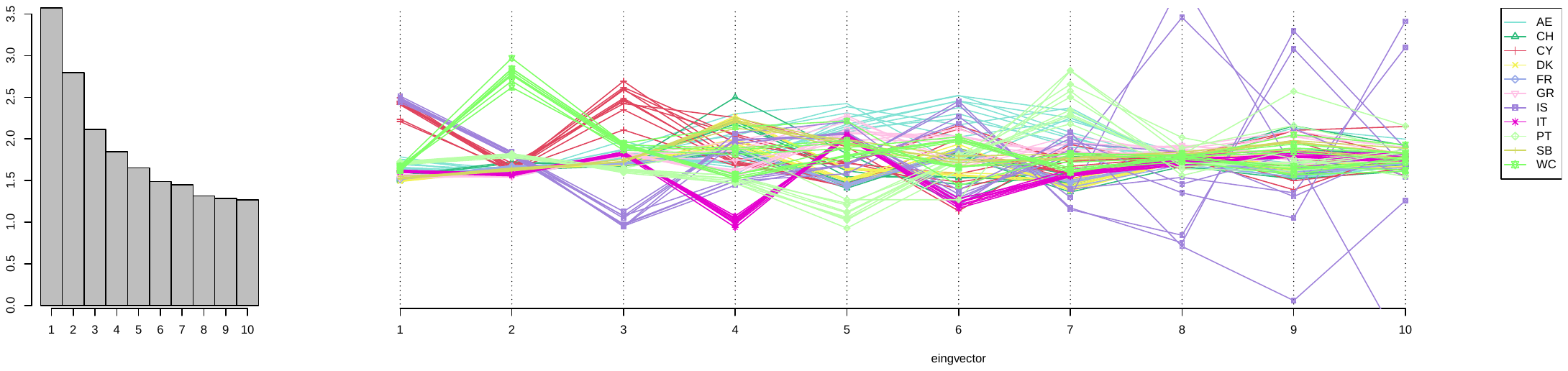

**Figure S2 –** Screeplot of the 10 fist axes of the PCA with all the individuals (left). Position of the individuals on the 10 first axes of the PCA (right).

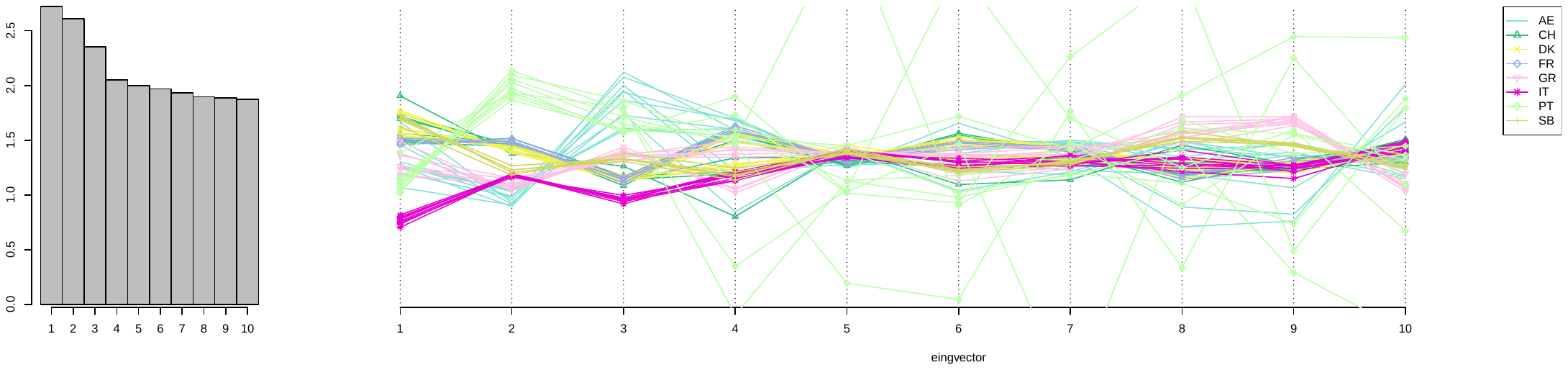

**Figure S3 -** Screeplot of the 10 fist axes of the PCA with 66 European individuals (left). Position of the individuals on the 10 first axes of the PCA (right).

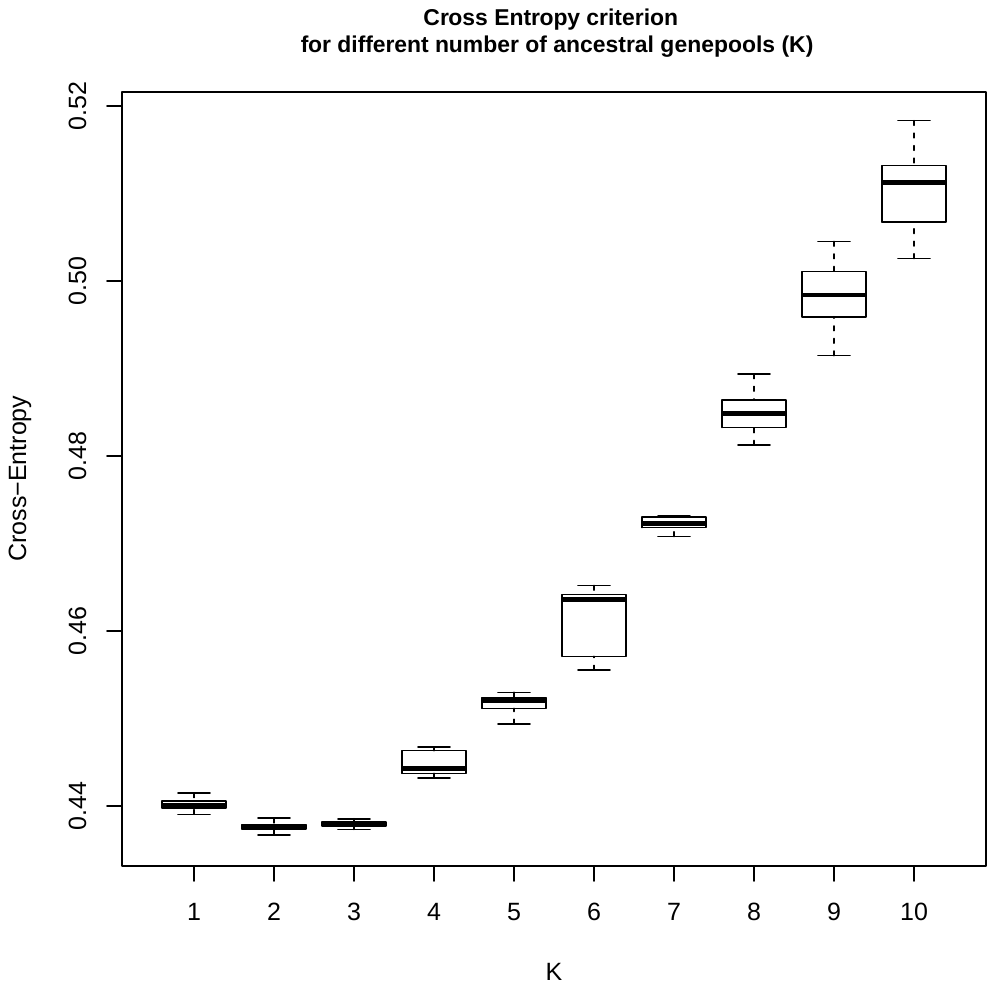

**Figure S4** – Values of the cross-entropy criterion for 25 sNMF runs per K. The number of clusters ranged from 1 to 10.

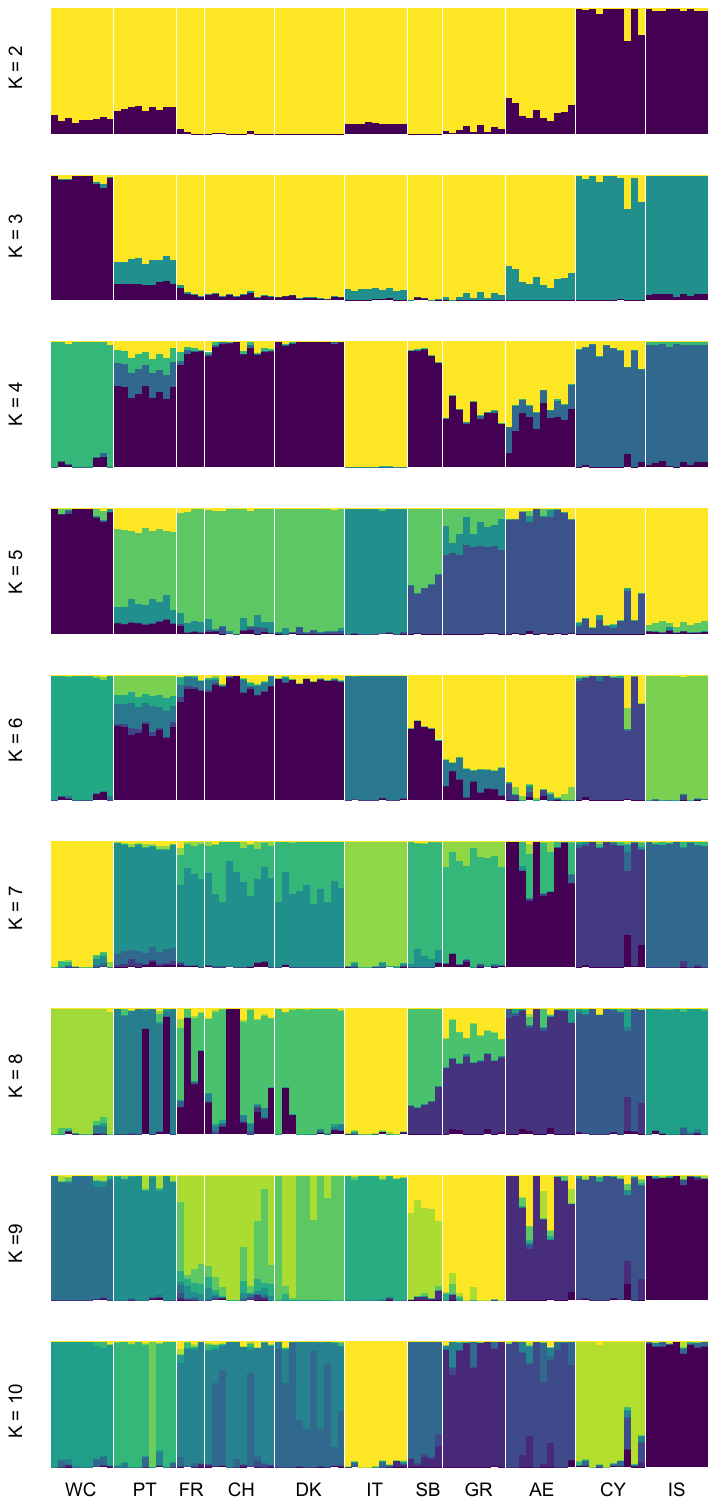

**Figure S5** – Individual ancestry estimated by sNMF for K ranging from 2 to 10. Each vertical bar represents one individual, and the colors represent the relative contributions of each genetic lineage K.

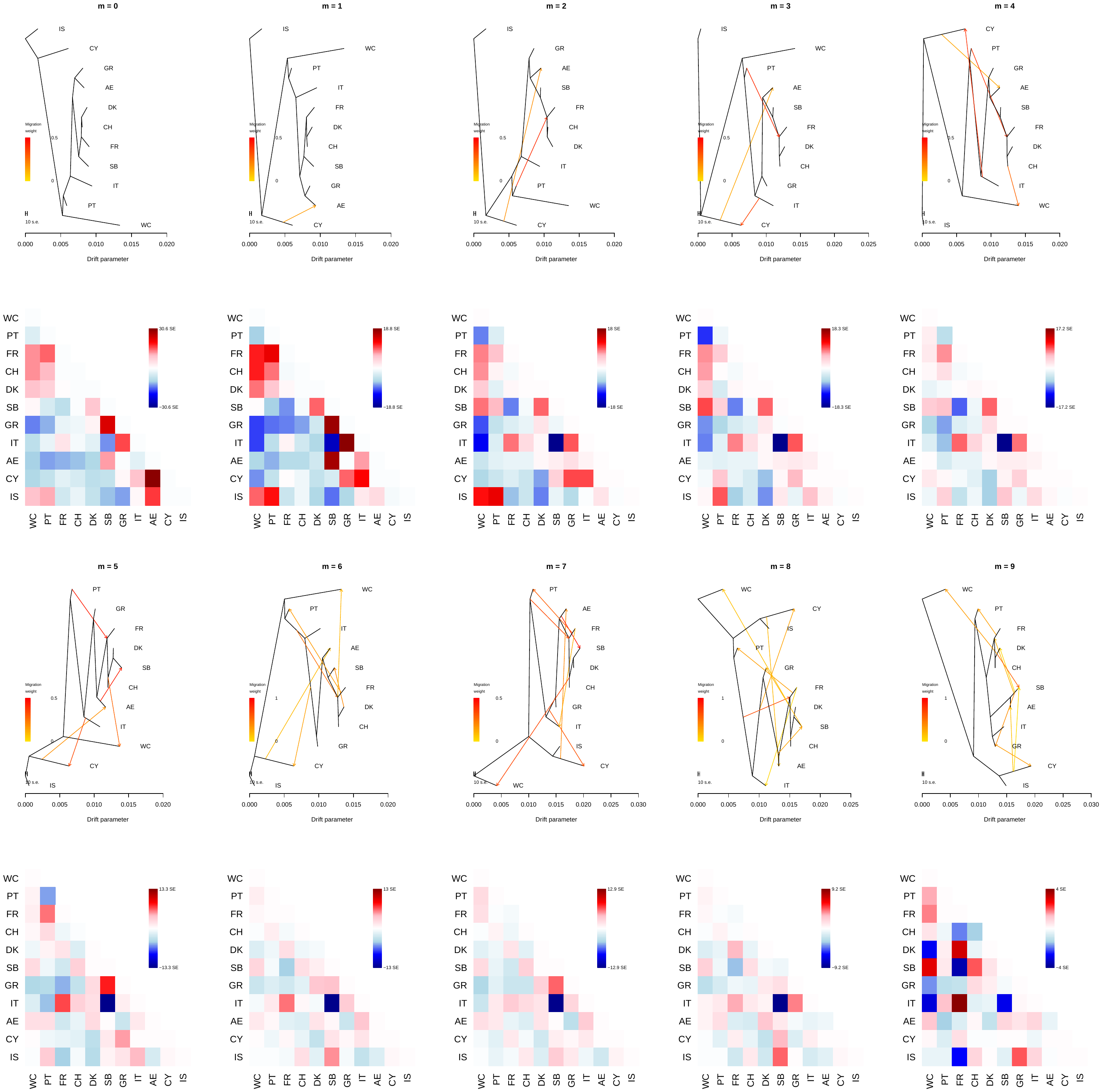

**Figure S6** – Results from TreeMix for 0 to 9 migration events. Highest likelihood runs are depicted, with the corresponding matrix of standard errors.

*Haplotype sharing*

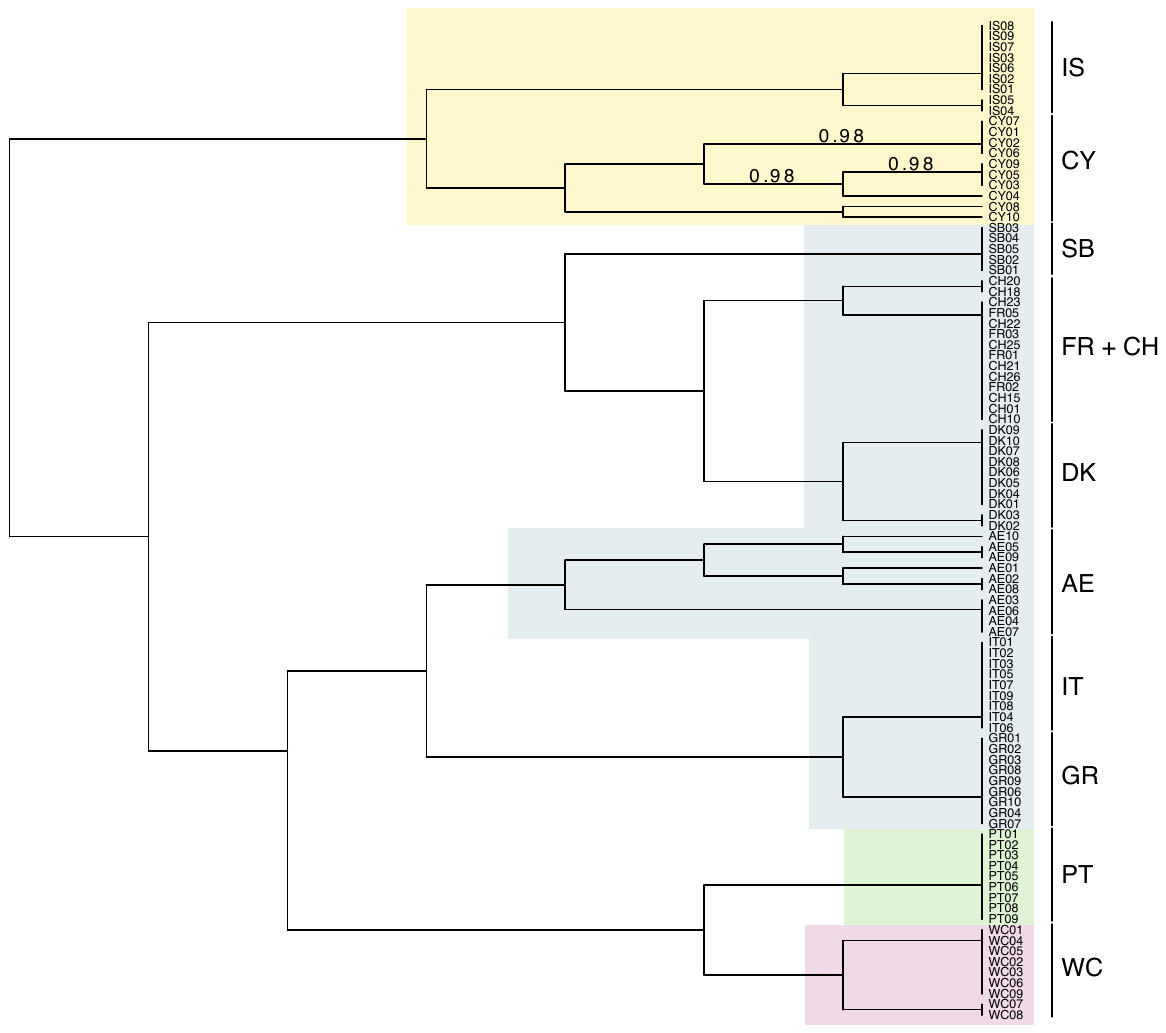

**Figure S7** – Dendrogram of the individuals from FineStrcture. Sampled populations were monophyletic, except for Swiss (CH) and French (FR) individuals. Background colors are consistent with the lineages identified in fig. 1. Grouping is based on similarity and the tree is not based on any model of population differentiation (*Lawson et al. 2012*).

**Modeling the History of European barn owl**

*Maximum-likelihood demographic inference*

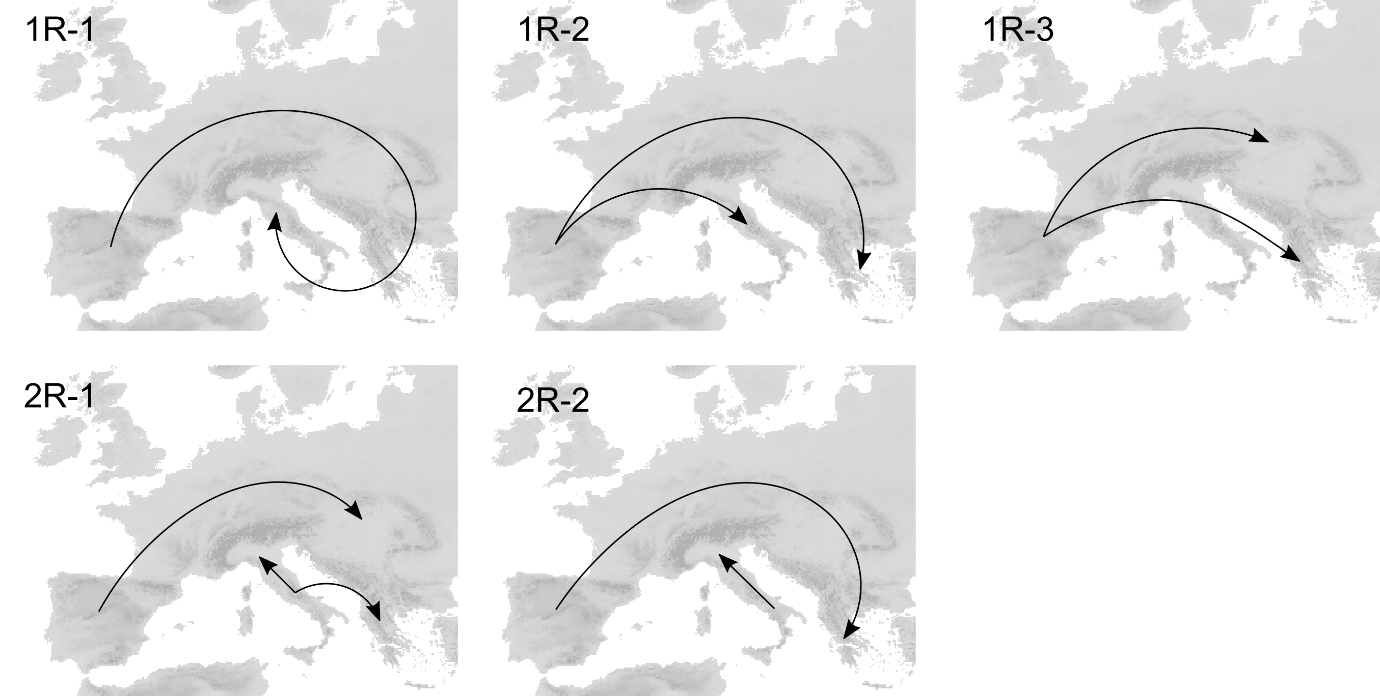

**Figure S8** – Schematic representation of the recolonization of Europe for the five scenarios modeled in *fastsimcoal2*.

**Table S2** – Parameter ranges and point estimate inferred for the demographic models tested with *fastsimcoal2*. Model “2R-1” – identified as the best fitting model – is given in Sup. Table 7. All range distributions were uniform except for bottlenecks (*) which were log-uniform. When a parameter was absent in a model, the case was left blank.

|  |  | **One Refugium** | | | **Two Refugia** |
| --- | --- | --- | --- | --- | --- |
| **Parameter** | **Ranges** | **1R -1** | **1R-2** | **1-R3** | **2R-2** |
| *Current Population Sizes (haploid)* | |  |  |  |  |
| PT | 10000 - 4e5 | 11713 | 11637 | 10218 | 386028 |
| CH | 1000 - 3.5e5 | 1009 | 194765 | 239033 | 261548 |
| GR | 1000 - 3.5e5 | 1031 | 1019 | 1037 | 4267 |
| IT | 1000 - 3.5e5 | 1022 | 5272 | 4692 | 296226 |
| *Ancestral Population Sizes (haploid)* | |  |  |  |  |
| PT before glac | 10000 - 4e5 | - | - | - | 244935 |
| IT before glac | 1000 - 3.5e5 | - | - | - | 115633 |
| PT during glac | 0.01 - 0.5 * | - | - | - | 654486 |
| IT during glac | 0.01 - 0.5 * | - | - | - | 83910 |
| *Times of Divergence (generations)* | |  |  |  |  |
| Pre-glacial split PT-IT | 8000 - 15000 | - | - | - | 8137 |
| T1 |  | 5954 | 5946 | 5873 | 5830 |
| T2 |  | 5937 | 5665 | 5853 | 0 |
| T3 |  | 3956 | 1453 | 1331 | - |
| *Current Migration (flow is backwards in time)* | | |  |  |  |
| CH → PT | 0 - 0.05 | 0.0074 | 0.0019 | 0.0001 | 0.0259 |
| PT → CH | 0 - 0.05 | 0.0029 | 0.0027 | 0.0037 | 0.0177 |
| GR → CH | 0 - 0.05 | 0.0212 | 0.0360 | 0.0172 | 0.0135 |
| CH → GR | 0 - 0.05 | 0.0038 | 0.0412 | 0.0182 | 0.0182 |
| IT → GR | 0 - 0.05 | 0.0164 | 0.0019 | 0.0016 | 0.0342 |
| GR → IT | 0 - 0.05 | 0.0005 | 0.0031 | 0.0057 | 0.0130 |
| IT → CH | 0 - 0.05 | 0.0013 | 0.0001 | 0.0001 | 0.0341 |
| CH → IT | 0 - 0.05 | 0.0002 | 0.0013 | 0.0001 | 0.0198 |
| *Older Migration* | | |  |  |  |
| CH → PT | 0 - 0.05 | 0.0007 | - | 0.0017 | 0.00001 |
| PT → CH | 0 - 0.05 | 0.0209 | - | 0.0249 | 0.0001 |
| GR → CH | 0 - 0.05 | 0.0047 | - | - | - |
| CH → GR | 0 - 0.05 | 0.0709 | - | - | - |
| *Instbot- instant founding population ** | | |  |  |  |
| Pre-glacial split PT-IT | 0.01 - 0.5 | - | - | - | 69907 |
| T1 | 0.01 - 0.5 | 84 | 189 | 99 | 12011 |
| T2 | 0.01 - 0.5 | 19 | 6750 | 3957 | 58 |
| T3 | 0.01 - 0.5 | 77 | 151 | 65 | - |

**Table S3 –** Likelihood and AIC of the demographic models tested with *fastsimcoal2*. Five main model topologies were tested with one (1R) or two (2R) glacial refugia. Models are sorted from best to worst according to the estimated likelihoods.

| **Model Name** | **Est. Lhood** | **Δ Lhood** | **AIC** | | **Δ AIC** |
| --- | --- | --- | --- | --- | --- |
| 2R-1 | -6519338 | 3052.89 | 30022708.26 | 0 | |
| 1R-1 | -6522237 | 5951.772 | 30036060.1 | 13351.85 | |
| 1R-3 | -6525492 | 9207.312 | 30051046.42 | 28338.16 | |
| 1R-2 | -6529137 | 12852.437 | 30067830.84 | 45122.58 | |
| 2R-2 | -6556984 | 40699.49 | 30196081.26 | 173372.99 | |

Est. Lhood – Maximum-likelihood estimated for the simulated SFS per demographic model; Δ Lhood – difference between the likelihood of the simulated and observed SFS; Δ AIC – delta AIC

**Table S4 –** Parameter point estimates and 95% confidence interval for the best demographic model: 2R -1. Times of divergence are in years calculated with a generation time of 3 years. Migration rates and number of individuals are given forward in time.

| **Parameter** | **Lower Limit CI** | **Point Estimate** | **Upper Limit CI** |
| --- | --- | --- | --- |
| *Current Population Sizes (haploid)* | | | |
| PT | 20405 | 44055 | 380159 |
| CH | 1006 | 3462 | 205409 |
| GR | 1007 | 1381 | 208303 |
| IT | 1039 | 1319 | 326428 |
| *Ancestral Population Sizes (haploid)* | | | |
| PT before glac | 10103 | 10859 | 73118 |
| IT before glac | 10775 | 189631 | 320792 |
| PT during glac | 700 | 6857 | 141005 |
| IT during glac | 36 | 62 | 116247 |
| *Times of Divergence* | | | |
| Pre-glacial split PT-IT | 24206 | 68907 | 90015 |
| T1 | 1719 | 5095 | 16007 |
| T2 | 1622 | 5717 | 17523 |
| *Current Migration (forward 2Nm)* | | | |
| CH → PT | 46.28 | 90.59 | 1413.66 |
| PT → CH | 2.65 | 8.05 | 62.01 |
| GR → CH | 0.13 | 41.66 | 156.06 |
| CH → GR | 0.11 | 1.26 | 95.91 |
| IT → GR | 0.21 | 2.88 | 135.06 |
| GR → IT | 0.16 | 3.89 | 60.28 |
| IT → CH | 0.17 | 26.94 | 157.00 |
| CH → IT | 0.20 | 0.02 | 37.80 |
| *Instbot- instant founding population ** | | | |
| Pre-glacial split PT-IT | 66 | 80 | 73468 |
| T1 | 18 | 119 | 16216 |
| T2 | 23 | 29 | 19330 |

*Niche modeling*

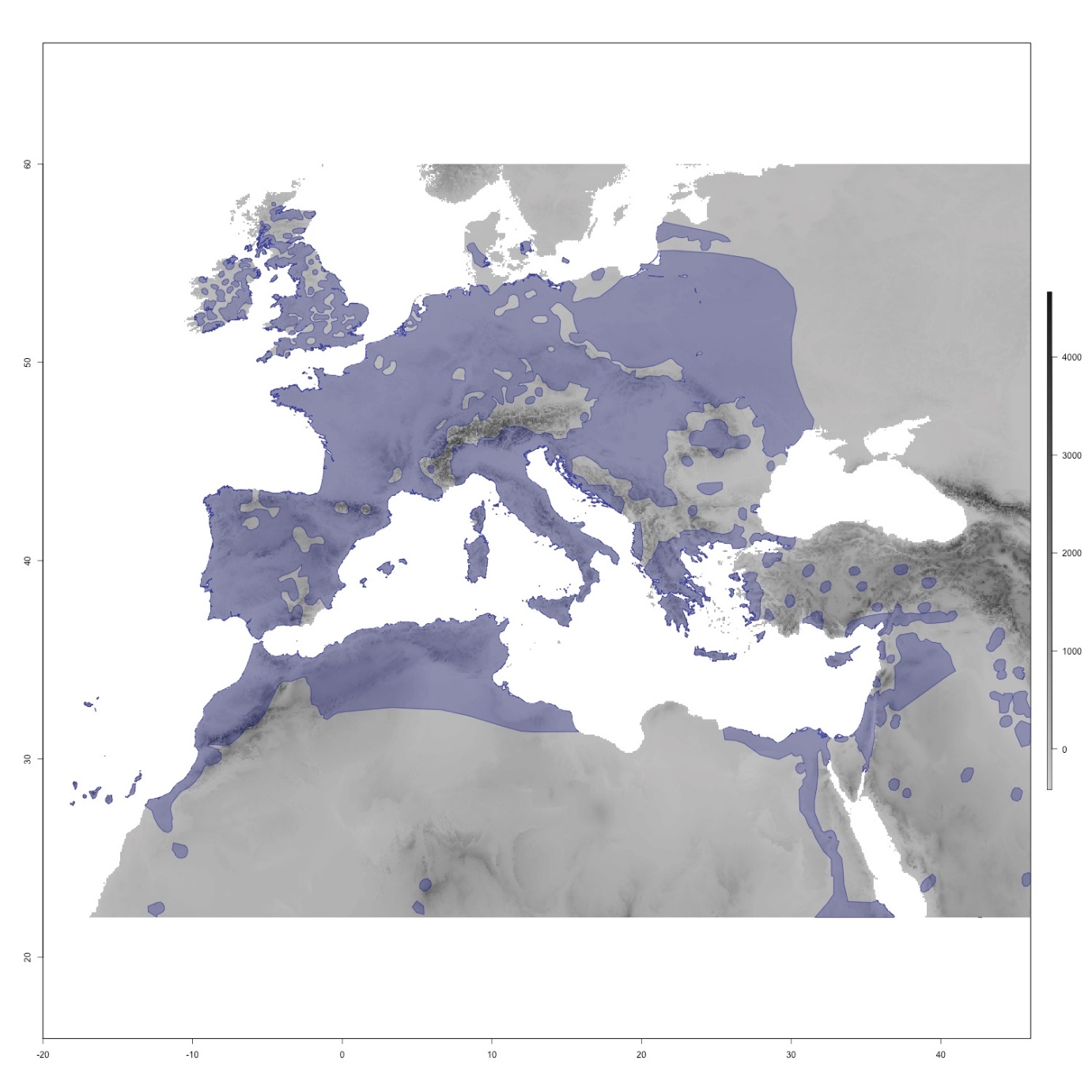

**Figure S9** – The map in grey depicts the area considered for producing the Species Distribution Model (SDM) for the barn owl. Shading is relative to the altitude. The current distribution of barn owls is plotted atop the map in purple (data from IUCN: BirdLife International 2019). Random presence points were extracted within this distribution for the SDM.

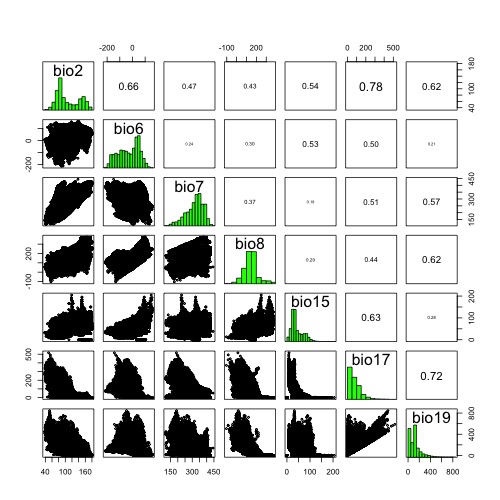

**Figure S10** – Pairwise correlation between climatic variables retained to produce the SDM. Only variables correlated at less than 0.8 were kept in the models, namely: Mean Diurnal Range (Bio2), Min Temperature of Coldest Month (Bio6), Temperature Annual Range (Bio7), Mean Temperature of Wettest Quarter (Bio8), Precipitation Seasonality (Bio15), Precipitation of Driest Quarter (Bio17) and Precipitation of Coldest Quarter (Bio19).

**Table S5** - Comparison of SDM model fit. AICc is reported for the multiple combinations of feature (linear, quadratic, hinge) and Beta multiplier (1 to 5).

|  | 1 | 2 | 3 | 4 | 5 |
| --- | --- | --- | --- | --- | --- |
| linear | 23953.4 | 23994.8 | 24037.1 | 24086.8 | 24125.8 |
| quadratic | **23950.8** | 24002.6 | 24042.7 | 24095.2 | 24148.5 |
| hinge | 24099.1 | 24185.4 | 24241.2 | 24304.4 | 24340.2 |

**Barriers and corridors**

*Migration Surface Estimate*

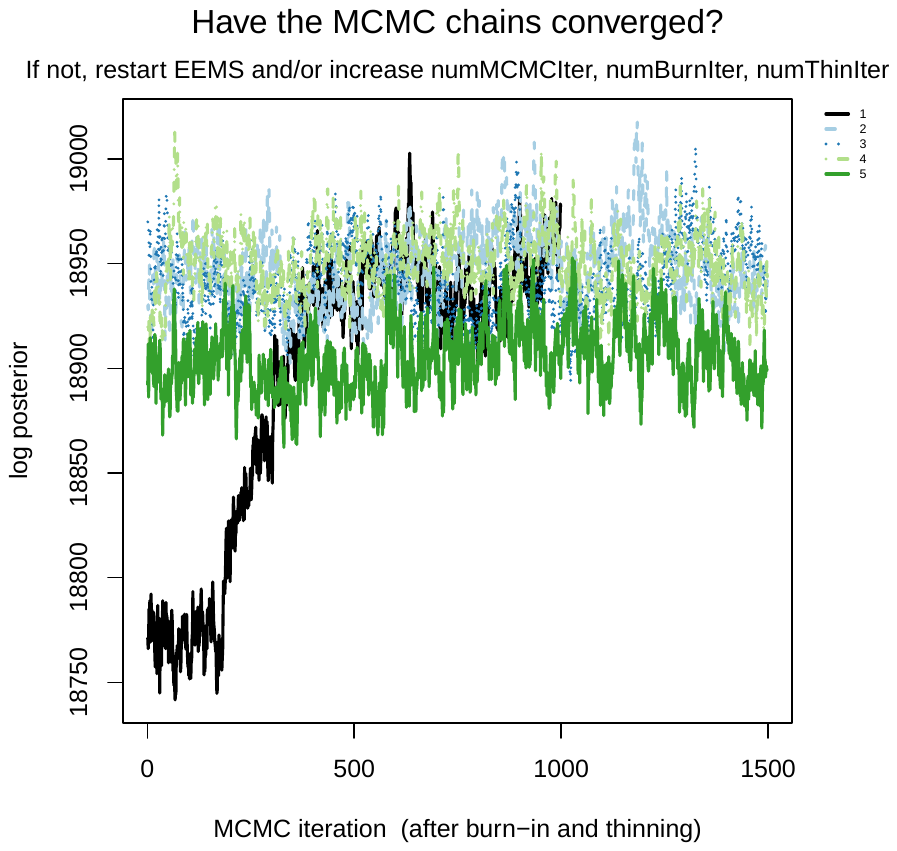

**Figure S11** - Convergence of MCMC chains for EEMS run in the Western Palearctic. Five independently seeded MCMC chains reach approximate convergence.

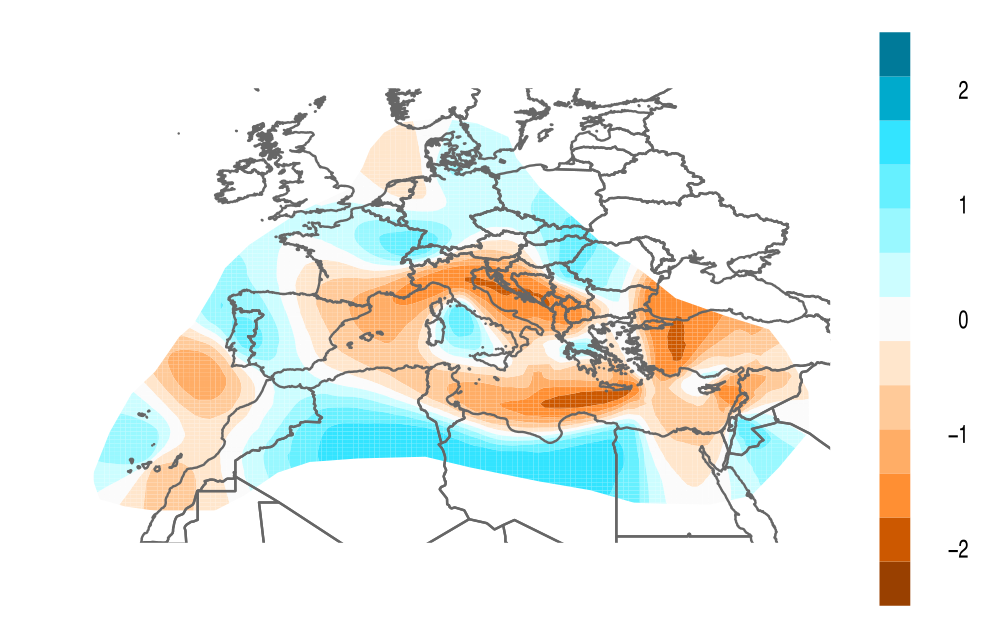

**Figure S12** – Effective migration surface (EEMS) of barn owls in the Western Palearctic. The map depicts relative barn owl migration in western Palearctic. Blue indicates a greater migration rate over the average; and orange a lower migration than average. Map shows the mean of 5 independent EEMS posterior migration rate estimates between 1000 demes.

*Isolation by distance*

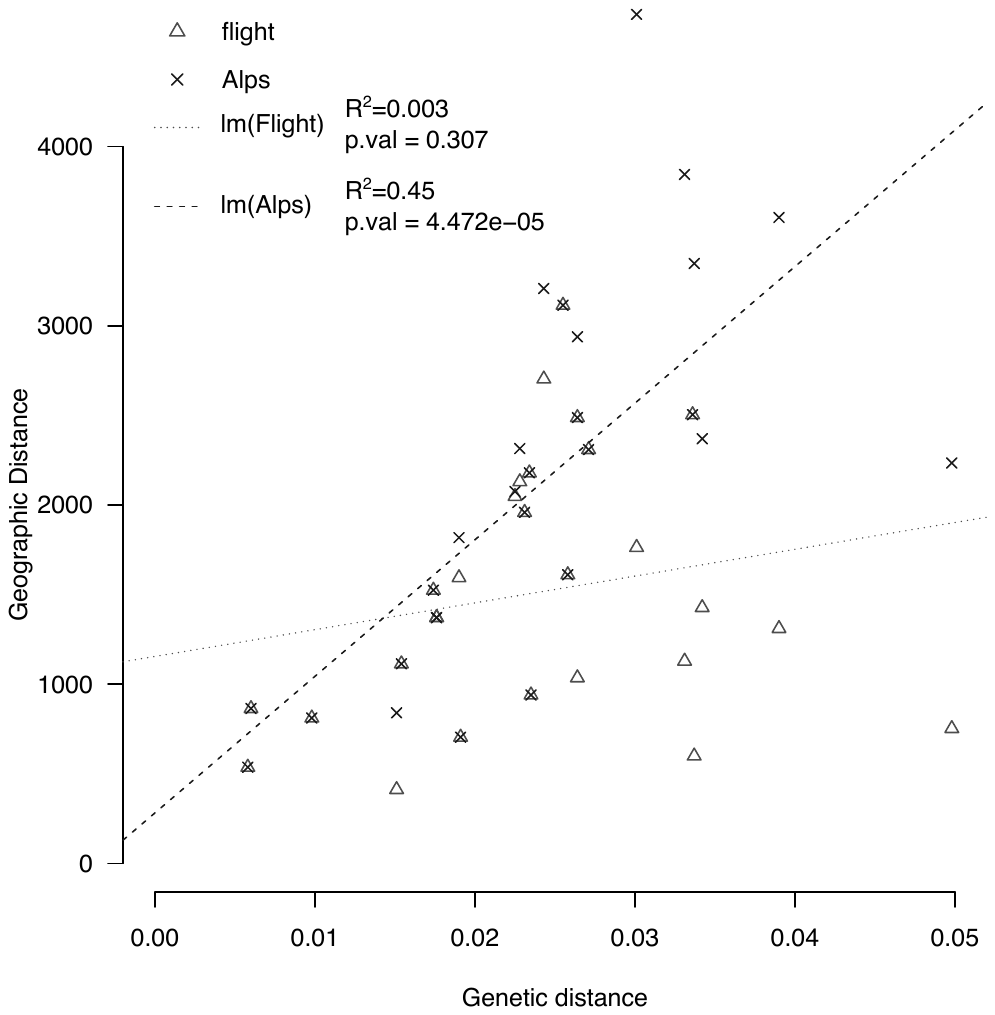

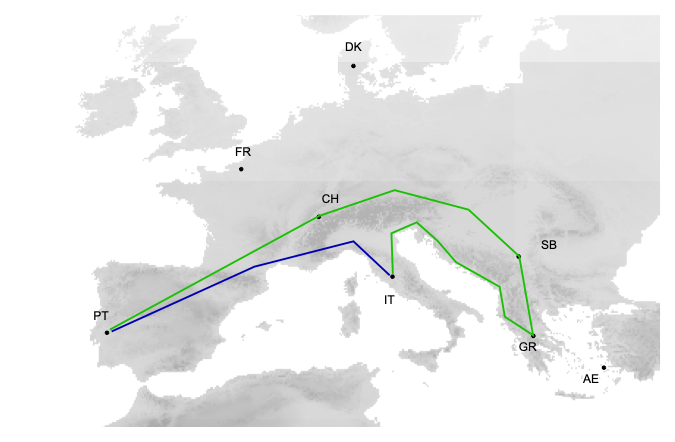

**Figure S13** – Isolation by distance in European barn owl. (a) Correlation of the pairwise genetic distance and two geographic distances between barn owl populations. Flight (Δ) refers to the straight-line flight distance over land between populations, while Alps (x) refers to distance between populations by considering the presence of the Alps. Lines depict the linear regression of both comparisons. R^2^ and p-values of each model are reported in the legend. (b) Alternative distance models. The example illustrates the shortest overland distance (blue), and distance around the Alps (green) between the populations from Italy (IT) and the Iberian Peninsula (PT).
